# Single nucleus analysis of the chromatin landscape in mouse forebrain development

**DOI:** 10.1101/159137

**Authors:** Sebastian Preissl, Rongxin Fang, Yuan Zhao, Ramya Raviram, Yanxiao Zhang, Brandon C. Sos, Hui Huang, David U. Gorkin, Veena Afzal, Diane E. Dickel, Samantha Kuan, Axel Visel, Len A. Pennacchio, Kun Zhang, Bing Ren

## Abstract

Genome-wide analysis of chromatin accessibility in primary tissues has uncovered millions of candidate regulatory sequences in the human and mouse genomes^1–4^. However, the heterogeneity of biological samples used in previous studies has prevented a precise understanding of the dynamic chromatin landscape in specific cell types. Here, we show that analysis of the transposase-accessible-chromatin in single nuclei isolated from frozen tissue samples can resolve cellular heterogeneity and delineate transcriptional regulatory sequences in the constituent cell types. Our strategy is based on a combinatorial barcoding assisted single cell assay for transposase-accessible chromatin^5^ and is optimized for nuclei from flash-frozen primary tissue samples (snATAC-seq). We used this method to examine the mouse forebrain at seven development stages and in adults. From snATAC-seq profiles of more than 15,000 high quality nuclei, we identify 20 distinct cell populations corresponding to major neuronal and non-neuronal cell-types in foetal and adult forebrains. We further define cell-type specific *cis* regulatory sequences and infer potential master transcriptional regulators of each cell population. Our results demonstrate the feasibility of a general approach for identifying cell-type-specific *cis* regulatory sequences in heterogeneous tissue samples, and provide a rich resource for understanding forebrain development in mammals.

## MAIN

A significant fraction of the non-coding DNA in the mammalian genome encodes transcriptional regulatory elements that play fundamental roles in mammalian development and human disease^3,6^. Identification of these sequences and characterizing their dynamic activities in specific cell types is a major goal in biology. Analysis of chromatin accessibility in primary tissues using assays such as DNase-seq^2,4^ and ATAC-seq^7,8^ has led to annotation of millions of candidate *cis* regulatory elements in the human and mouse genomes^1,3^. Yet, the catalogue of current *cis* regulatory elements, based primarily on analysis of bulk, heterogeneous biological samples, lacks precise information regarding cell-type-and developmental-stage-specific activities of each element. *In-vivo* lineage tracing using INTACT mouse models^8,9^ and isolation of particular cell types based on specific protein markers can address this limitation to some degree and in limited cell types^10,11^. But a more general strategy is necessary to study primary tissues from all stages of development in human as well as in other species.

In theory, single cell based chromatin accessibility studies can be used for unbiased identification of subpopulations in a heterogeneous population, and proof of principle has been reported using cultured mammalian cells and cryopreserved blood cell-types^5,12,13^. However, there has been no report that such approaches have been successfully used to dissect transcriptional regulatory landscapes in primary tissues. The main difficulty is that primary tissues are typically preserved either by formalin fixed paraffin embedding or flash freezing, conditions that are not amenable for isolating intact single cells. Here, we show that it is possible to isolate single nuclei from frozen tissues and assay chromatin accessibility in each nucleus in a massively parallel manner.

We adopted a combinatorial single cell ATAC-seq (scATAC-seq) strategy^5^ and optimized it for frozen tissue sections (Methods). Compared to the previous protocol^5^, key modifications were made to maximally preserve nuclei integrity during sample processing and optimize transposase-mediated fragmentation of chromatin in individual nuclei (Extended Data Fig. 1-3). We applied this improved protocol, hereafter referred to as snATAC-seq (<u>s</u>ingle <u>n</u>uclei ATAC-seq), to mouse forebrain in the 8-week-old adult mouse (P56) and in seven developmental stages at 1-day intervals starting from embryonic day 11.5 (E11.5) to birth (P0) (Fig.1a, b). Sequencing libraries were sequenced to almost saturation as indicated by a read duplication rate of 36-73% per sample (Extended Data Table 1). We filtered out low quality datasets using three stringent quality control criteria including read depth (Extended Data Fig. 3d), recovery rate of constitutively accessible promoters in each nucleus (Extended Data Fig. 3e), and signal-over-noise ratio estimated by fraction of reads in peak regions (Method; Extended Data Fig. 3f). In total, 15,767 high-quality snATAC-seq datasets were obtained. The median read depth per nucleus ranged from 9,275 – 18,397, median promoter coverage was 11.6%, and the median fraction of reads in peak regions was 22% (Extended Data Table 2, 3). Our protocol maintains the extraordinary scalability of combinatorial indexing, while featuring a ∼ 6 fold increase in read depth per nucleus (Extended Data Table 3). The high quality of the single nuclei chromatin accessibility maps was supported by a high concordance between the aggregate snATAC-seq data and bulk ATAC-seq data (R > 0.9), and high reproducibility between biological replicates (R > 0.91, Fig. 1c, Extended Data Fig. 4).

**Figure 1:**
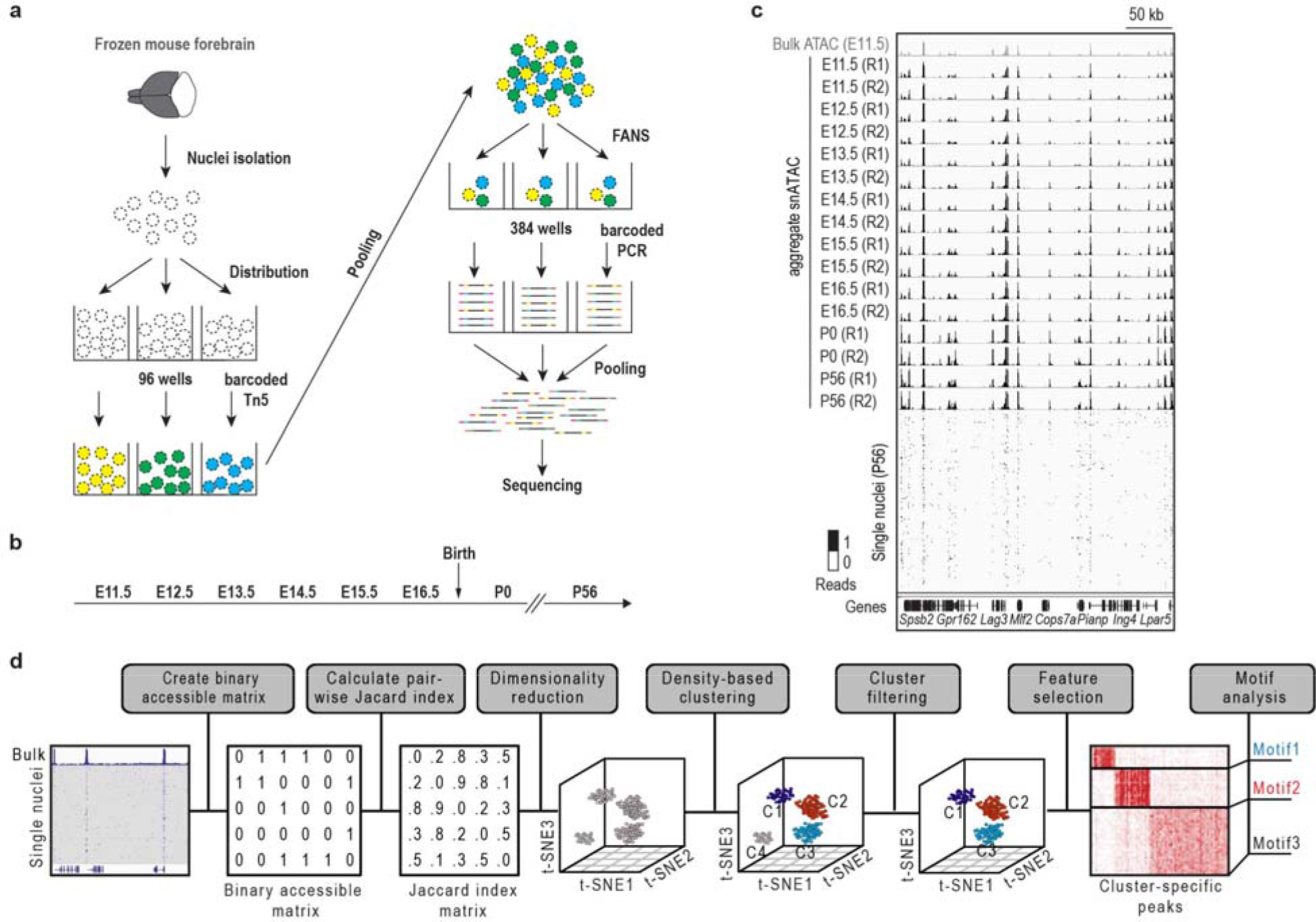
Experimental overview and computational analysis strategy for snATAC-seq. **a** Following nuclei isolation from frozen forebrain tissue biopsies, tagmentation of 4,500 permeabilized nuclei was carried out using barcoded Tn5 in 96 wells. After pooling, 25 nuclei were sorted in 384 wells and PCR-amplified to introduce second barcodes. FANS: Fluorescence assisted nuclei sorting. **b** Overview of investigated time points during mouse development. E: embryonic day; P: postnatal day; **c** Chromatin accessibility profiles of aggregate snATAC-seq (black tracks) agree with bulk ATAC-seq (grey, top track) and are consistent between biological replicates. **d** Framework of computational analysis of snATAC-seq data.

**Extended Data Figure 1:**
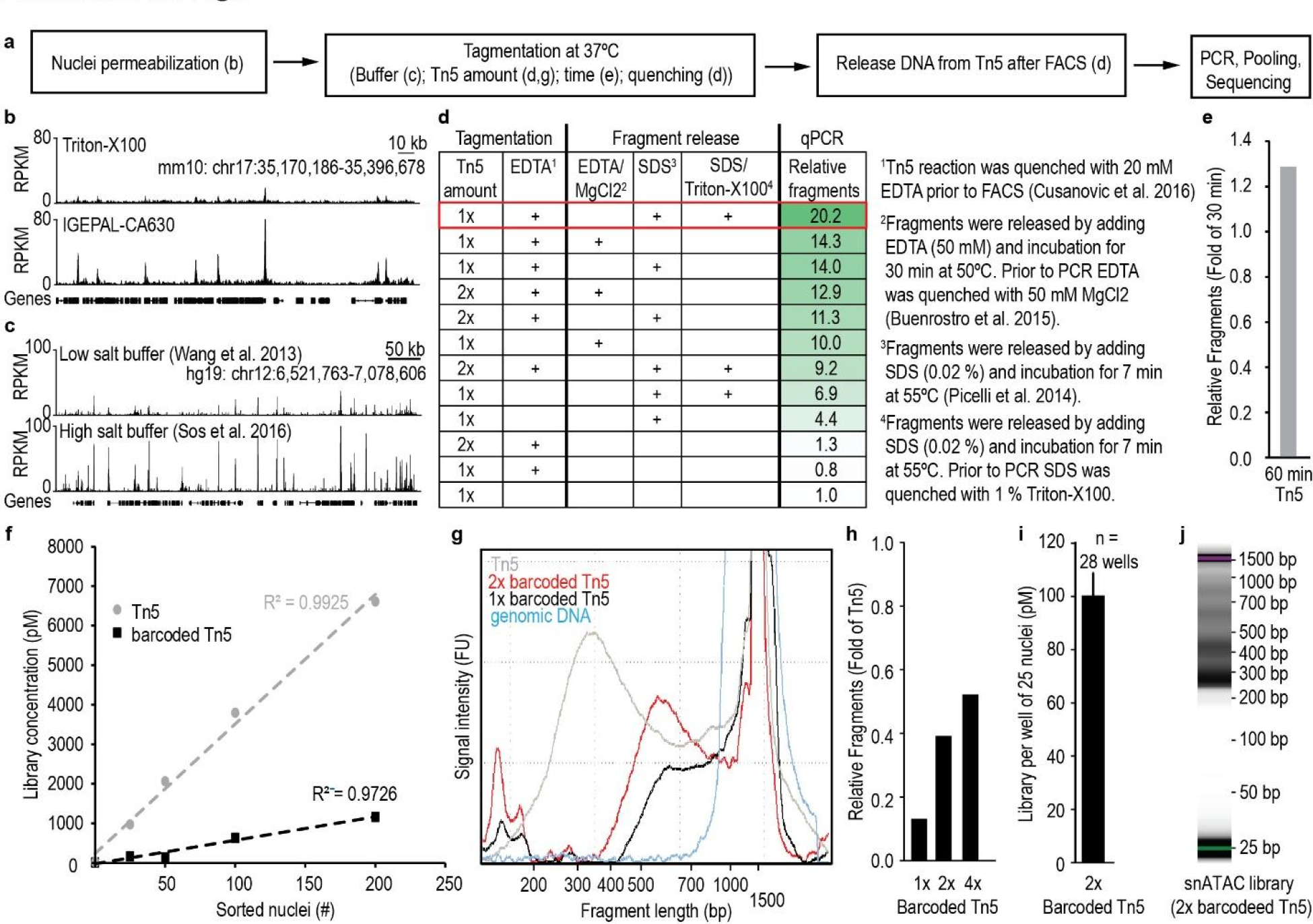
SnATAC-seq protocol optimization. **a** Overview of critical steps for the snATAC-seq procedure for nuclei from frozen tissues. **b** IGEPAL-CA630 but not Triton-X100 was sufficient for tagmentation of frozen tissues. **c** Tagmentation was facilitated by high salt concentrations in reaction buffer^47,48^. **d** Maximum amount of fragments per nucleus could be recovered when quenching Tn5 by EDTA prior to FACS and denaturation of Tn5 after FACS by SDS. Finally, SDS was quenched by Triton-X100 to allow efficient PCR amplification. **e** Increasing tagmentation time from 30 min to 60 min can result in more DNA fragments per nucleus. **f** Number of sorted nuclei was highly correlated with the final library concentration. Tn5 loaded with barcoded adapters showed less efficient tagmentation as compared to Tn5 without barcodes. Wells were amplified for 13 cycles, purified and libraries quantified by qPCR using standards with known molarity. **g** Tagmentation with barcoded Tn5 was less efficient and resulted in larger fragments than Tn5 (550 bp vs. 300 bp). Ratio for barcoded Tn5 was based on concentration of regular Tn5. **h** Doubling the concentration of barcoded Tn5 significantly increased the number of fragments per nucleus by 3 fold. Further increase resulted only in minor improvements. **i** Generated amount of library from 25 nuclei per well was reproducible between single wells. Each well was amplified for 11 cycles and quantified by qPCR. This output was used to calculate the number of required PCR cycles for snATAC-seq libraries to prevent overamplification (n = 28 wells, average ± SEM). **j** Size distribution of a successful snATAC-seq library from a mixture of E15.5 forebrain and GM12878 cells shows a nucleosomal pattern. SnATAC-seq was performed including all the optimization steps described above with barcoded Tn5 in 96 well format.

**Extended Data Figure 2:**
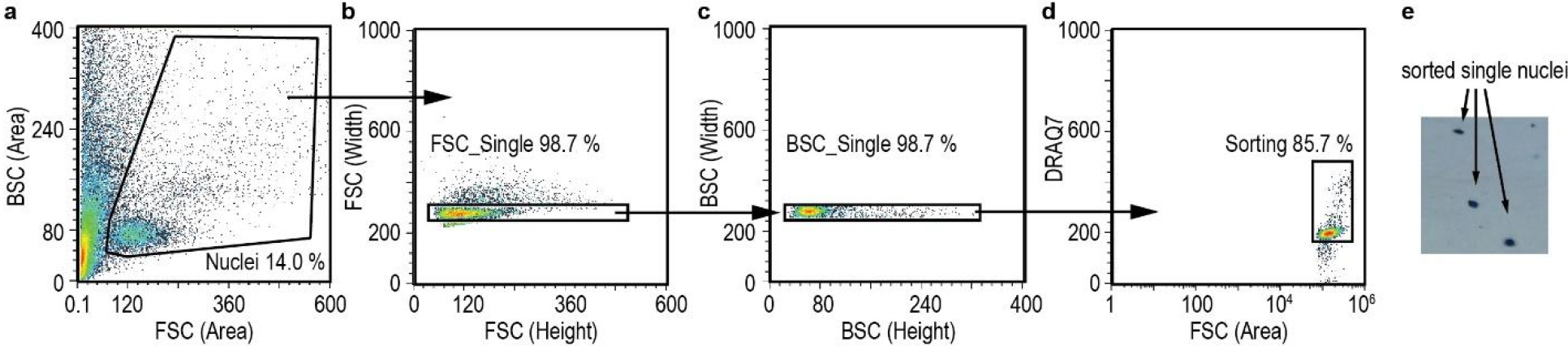
Sorting of single nuclei after tagmentation. **a-d** Density plots illustrating the gating strategy for single nuclei. First, big particles were identified (**a**), then duplicates were removed (**b**, **c**) and finally, nuclei were sorted based on high DRAQ7 signal (**d**), which stains DNA in nuclei. **e** Verification of single cell suspension after FACS was done with Trypan Blue staining under a microscope.

**Extended Data Figure 3:**
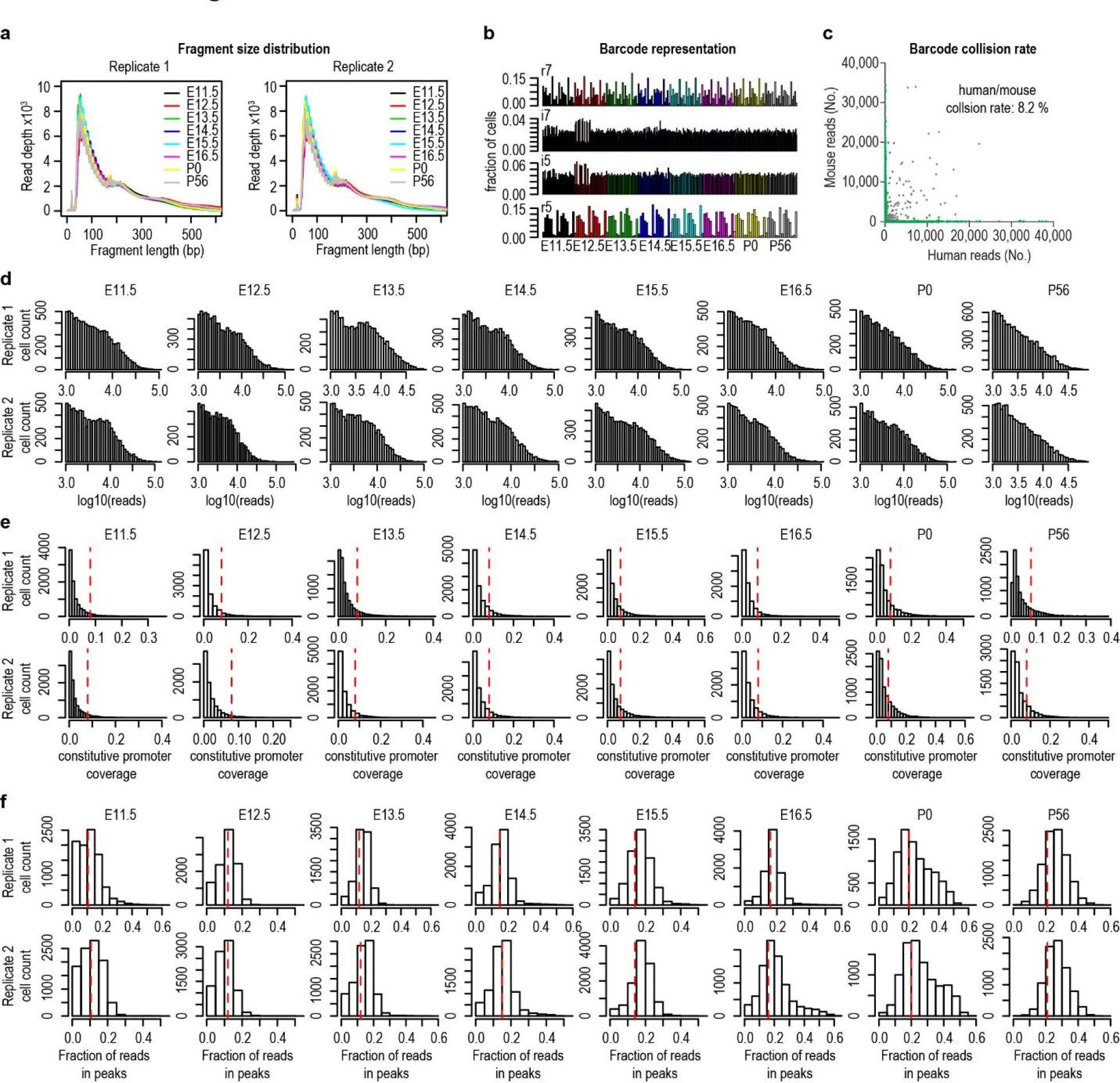
Overview of sequencing data and quality filtering for single cells. **a** Distribution of insert sizes between reads pairs derived from sequencing of snATAC-seq libraries indicates nucleosomal patterning. **b** Individual barcode representation in the final library shows variability between barcodes. **c** To assess the probability of two cells sharing the same cell barcode, single cell ATAC-seq was performed on a 1:1 mixture of human GM12878 cells and mouse E15.5 forebrain nuclei. A collision was indicated by <90% of all reads mapping to either the mouse genome (mm9) or the human genome (hg19). We identified 8.2% of these barcode collision events. **d**Read coverage per barcode combination after removal of potential barcodes with less than 1,000 reads. **e** Constitutive promoter coverage for each single cell. The red line indicates the constitutive promoter coverage in corresponding bulk ATAC-seq data sets from the same biological sample. Cells with less coverage than the bulk ATAC-seq data set were discarded. **f** Fraction of reads falling into peaks for each single cell. The red line indicates fraction of reads in peak regions in corresponding bulk ATAC-seq data sets from the same biological sample. Cells with lower reads in peak ratios coverage than the bulk ATAC-seq data set were discarded from downstream analysis. Bulk ATAC-seq data for E11.5-P0 were reanalysed (Zhao et al. manuscript in preparation)

**Extended Data Figure 4:**
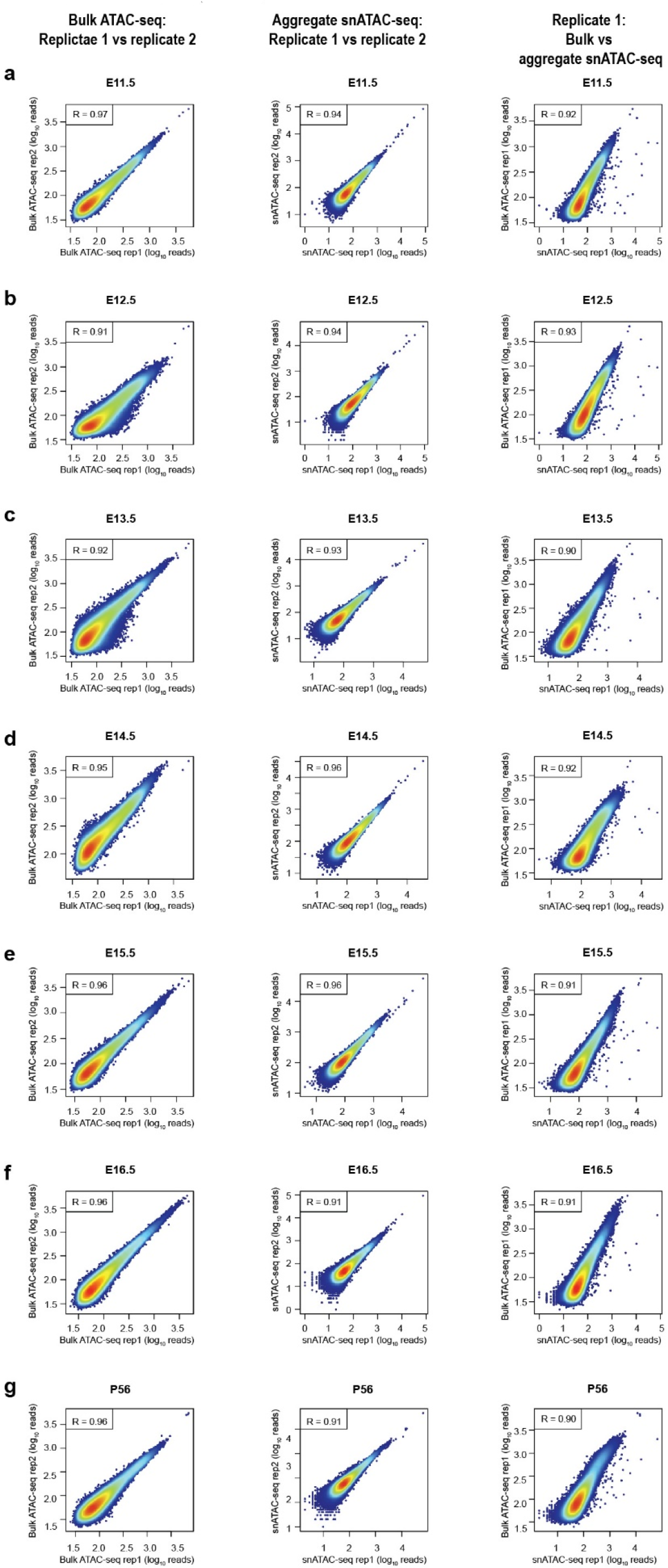
Pearson correlation plots of bulk and aggregate single-nuclei ATAC-seq data sets. Pearson correlation of chromatin accessibility profiles from two biological replicates derived from bulk ATAC-seq (left column) and from aggregate snATAC-seq after aggregating single cell profiles (middle column). In the right column the correlation between bulk ATAC-seq and aggregate snATAC-seq are displayed for biological replicate 1. Data are displayed from forebrain tissues from following time points: **a** E11.5, **b** E12.5 **c** E13.5 **d** E14.5 **e** E15.5 **f** E16.5 **g** P0 **h** P56. Bulk ATAC-seq data for E11.5-P0 were reanalysed (Zhao et al. manuscript in preparation).

**Extended Data Table 1:**
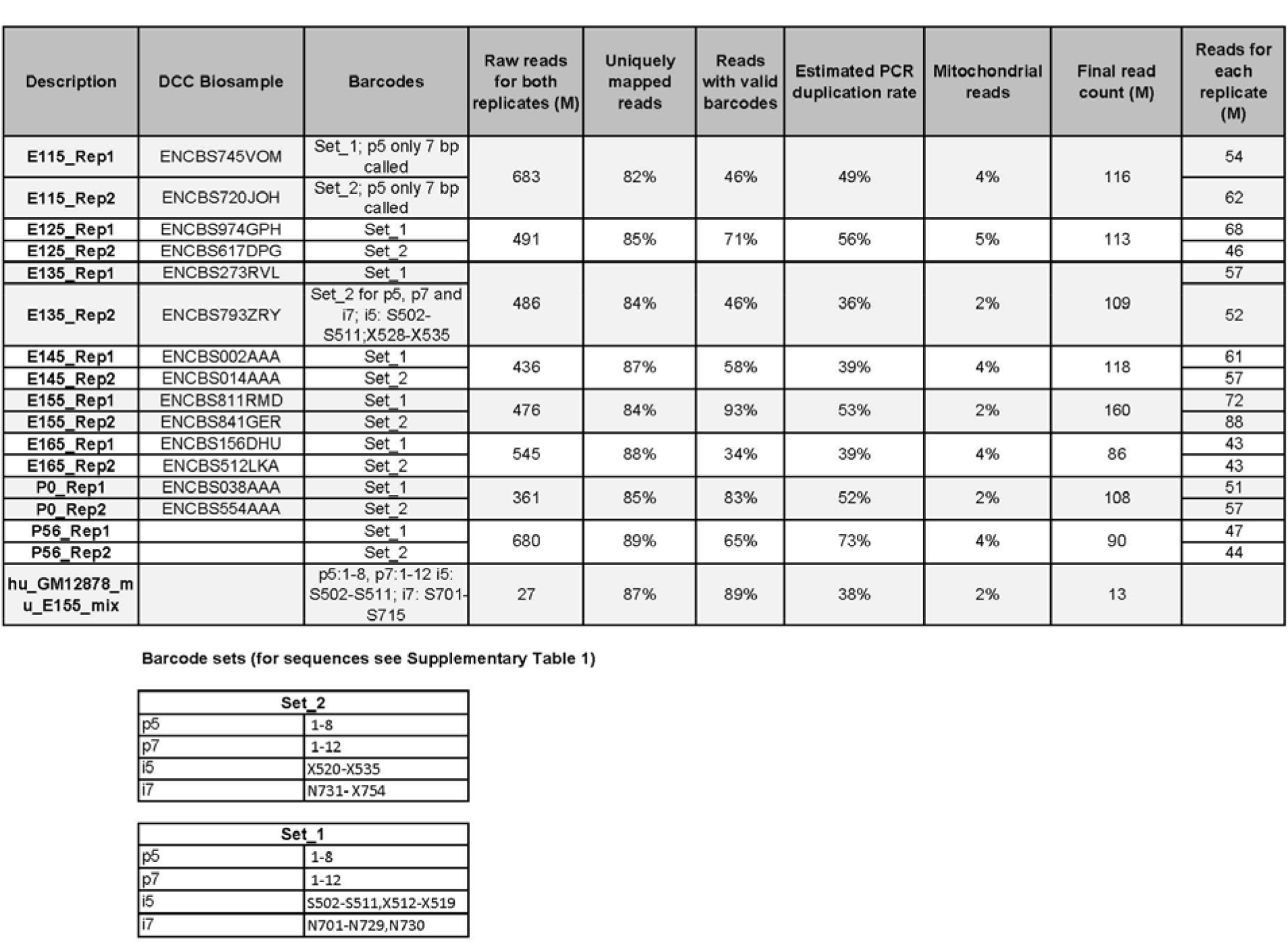
Sequencing statistics for single nuclei ATAC-seq libraries. General overview of sequencing for single cell ATAC-seq libraries including PCR duplication rates and fraction of mitochondrial reads. Please note that paired end reads were treated as separate reads. Replicate 1 and 2 were sequenced together and single cell datasets were assigned based on replicate specific barcode combinations (Set_1 or Set_2). One exception was E13.5 where replicate 1 and 2 were sequenced on separate lanes. Please note that for E11.5 7 out of 8 bp were detected for the p5 barcode. M: million

**Extended Data Table 2:** Overview of single nuclei ATAC-seq data after filtering out low quality cells. Overview of cells that pass quality control and general properties of data sets including promoter coverage and fraction of reads in peaks.

| Sample | Cells pass QC (#) | Reads per cell (median) | Promoter coverage (median) | Fraction of reads in peaks (median) |
| --- | --- | --- | --- | --- |
| E11.5_Rep1 | 528 | 16,023 | 10.6% | 17.5% |
| E11.5_Rep2 | 685 | 16,499 | 11.1% | 17.5% |
| E12.5_Rep1 | 781 | 18,397 | 11.2% | 15.7% |
| E12.5_Rep2 | 303 | 17,945 | 10.3% | 14.9% |
| E13.5_Rep1 | 651 | 16,891 | 11.8% | 18.4% |
| E13.5_Rep2 | 646 | 14,669 | 11.1% | 18.7% |
| E14.5_Rep1 | 976 | 14,440 | 11.3% | 19.8% |
| E14.5_Rep2 | 905 | 14,489 | 11.5% | 19.2% |
| E15.5_Rep1 | 1,591 | 16,789 | 12.2% | 20.4% |
| E15.5_Rep2 | 2,235 | 17,840 | 12.9% | 20.5% |
| E16.5_Rep1 | 317 | 12,758 | 11.0% | 23.2% |
| E16.5_Rep2 | 738 | 10,460 | 11.4% | 29.3% |
| P0_Rep1 | 1,044 | 9,275 | 13.3% | 36.0% |
| P0_Rep2 | 1,333 | 8,703 | 12.6% | 37.4% |
| P56_Rep1 | 1,569 | 12,509 | 11.6% | 27.1% |
| P56_Rep2 | 1,465 | 12,689 | 11.5% | 26.4% |
| Average | 985 | 14,399 | 11.6% | 22.6% |
| Total | 15,767 |  |  |  |

**Extended Data Table 3:** Comparison of single nuclei ATAC-seq with previously published initial single cell/nuclei ATAC-seq studies. Table illustrating several characteristics of single nuclei/cell ATAC_seq library. n.r. not reported

| Parameter | this study | Cusanovich et al. 2015<sup>5</sup> | Buenrostro et al. 2015<sup>12</sup> |
| --- | --- | --- | --- |
| Single cell strategy | Combinatorial indexing | Combinatorial indexing | Microfluidics |
| Sample type | Frozen tissues | Cell lines | Cell lines |
| Total cells/nuclei (per experiment) | 15,676 (303-2,235) | 15,814 (533-1,459) | 1886 (n.r.) |
| Reads per cell/nucleus (median) | 9,375-18,397 | 1,390-3,094 | 73,000 (average) |
| Fraction of reads in peaks | 15.7%-37.4% | 41-59 % | >15% |
| Promoter coverage per cell/nucleus | 11.60% | n.r. | 9.40% |

The snATAC-seq profiles from each forebrain tissue arise from a mixture of distinct cell types. We reasoned that cells of the same type should share higher similarity in the open chromatin profiles than cells from different cell types. Based on this assumption, we developed a computational framework to uncover distinct cell types from the snATAC-seq datasets without prior knowledge of cell types in the tissue. Specifically, we first determined the open chromatin regions from the bulk ATAC-seq profiles of mouse forebrain tissue in seven fetal development time points and in adults, resulting in a total of 154,364 combined open chromatin regions that were detectable in one or more stages (Fig. 1d, Methods, Supplementary Table 1; Zhao et al. manuscript in preparation). Next, we constructed a binary matrix of open chromatin regions, using 0 and 1 to indicate absence and presence of a read at each open chromatin region, respectively, in each nucleus (Fig. 1d). Third, we calculated the pairwise similarity between cells using a Jaccard index. After applying a non-linear dimensionality reduction method, t-SNE^14^, the Jaccard index matrix was projected to a low-dimension space to reveal cell clusters (Fig. 1d)^15^. Finally, we filtered out any cluster with abnormal sequencing depth or in-group similarity compared to other clusters^5^. We performed the clustering using the 140,103 distal elements (outside 2 kb upstream of refSeq transcription start site), since previous studies have shown highly cell type-specific chromatin accessibility profiles at enhancer regions^16^, and that such sequences were more effective at classifying cell types than promoter or transcriptomics data^12^ (Extended Data Fig. 5a, b).

**Extended Data Figure 5:**
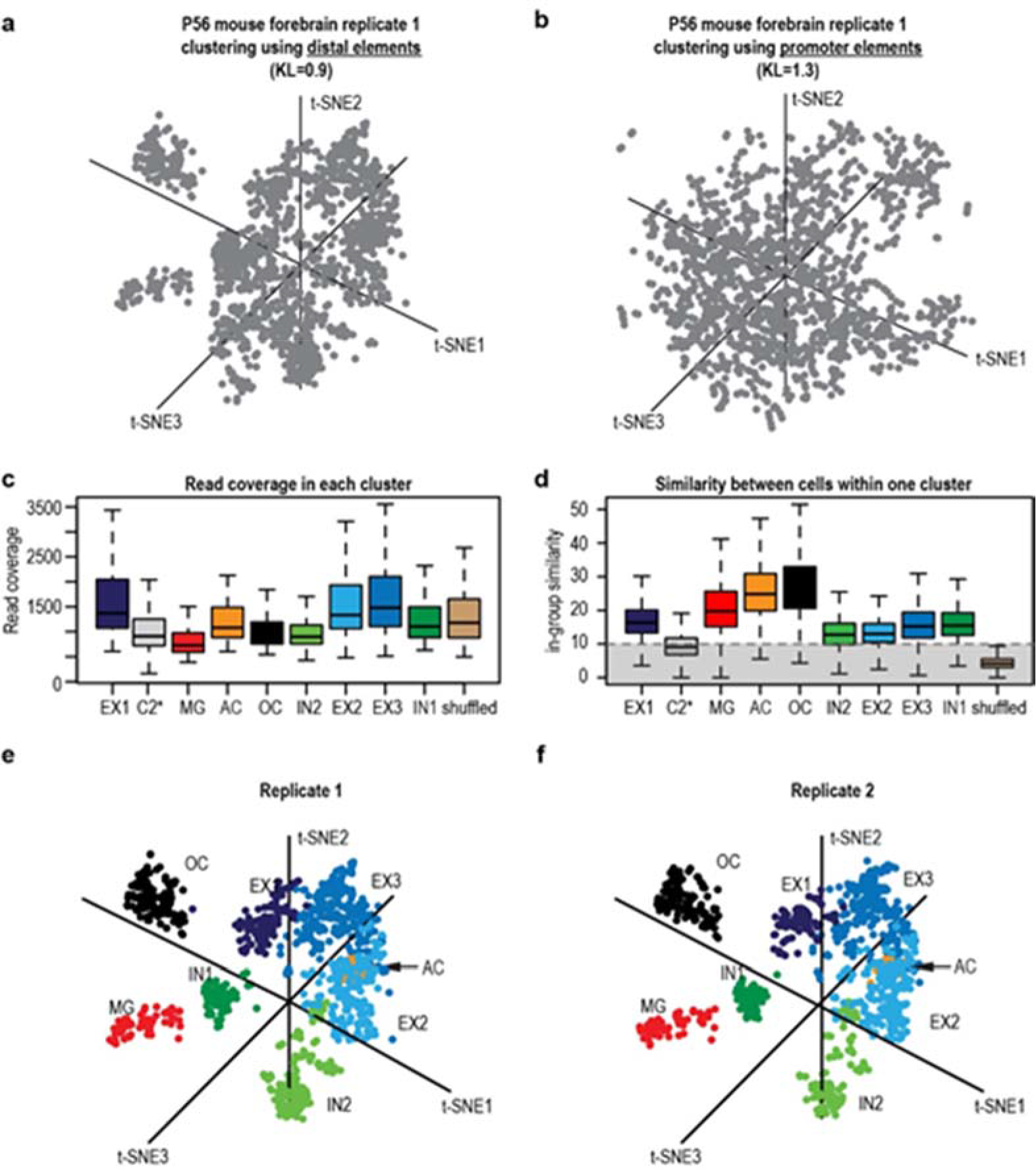
Clustering strategies, quality control of clusters and clustering result for individual replicates in adult forebrain. **a, b** T-SNE visualization of clustering using **a** distal elements (regions outside 2 kb of refSeq transcriptional start sites) or **b** promoter regions (KL: Kullback-Leibler divergence reported by t-SNE).**c** Box plot of read coverage for each cluster. **d** Box plot of similarity analysis between two any two given cells in a cluster. Cluster C2 was discarded before downstream analysis due to low its intra-group similarity (median < 10). As a negative control, randomly shuffled cells were included in the analysis displaying exceptionally low in-group similarity. **e, f** T-SNE visualization of single cells from **e** replicate 1 and **f** replicate 2. The projection and color coding is the same as in Fig. 2d.

To demonstrate the effectiveness of the above approach in uncovering cell type-specific chromatin landscapes from heterogeneous tissue samples, we first analysed the 3,033 snATAC-seq profiles obtained from the adult forebrain (Extended Data Table 2). As negative controls, we included 200 “shuffled” nuclear profiles (Extended data Fig.5c, d, Methods). Initially, nine discrete cell populations, in addition to the group representing shuffled cells, were uncovered (Extended Data Fig. 5c, d). The cluster C2 (including 946 nuclei), like the shuffled cells, exhibited significantly lower intra-group similarity than other clusters, and thus were not included in further downstream analysis (Extended Data Fig. 5c, d). None of the t-SNE dimensions was correlated with read depth (R < 0.3 for all dimensions) and the clustering results were reproducible between two biological replicates (Extended data Fig. 5e, f). To categorize the final eight cell populations, we analysed transposase accessible chromatin at known cell type-specific gene loci and compared it to published data from sorted excitatory neurons^8^, GABAergic neurons^9^, microglia^17^ and NeuN negative nuclei which mostly comprise non-neuronal cells including astrocytes and oligodendrocytes^18^ (Fig. 2b, Extended Data Fig. 6a-c). Three cell populations and the sorted excitatory neurons showed high accessibility at the gene locus of the terminal neuronal differentiation factor *Neurod6* and other excitatory neuron-specific genes^19^ (Fig. 2b, Extended Data Fig. 7a). Likewise, two cell clusters and the sorted GABAergic neurons showed similar accessibility at the GABA synthesis enzyme *Gad1* locus (Fig. 2b, Extended Data Fig. 7b)^20^. Using this strategy we were able to identify an astrocyte subpopulation according to the accessibility at the *Apoe* locus and other known astroglia markers^21^, an oligodendrocyte subpopulation based on the myelin-associated glycoprotein *Mog* and other oligodendrocyte marker genes^22^, and a microglia subpopulation based on complement factors including *C1qb* (Fig. 2b, Extended Data Fig. 7c-e)^17^. The categorization of cell groups was further confirmed by hierarchical clustering, with one remarkable exception that the inhibitory neuron cluster 2 (IN2) clustered with excitatory neurons (Fig. 2c). According to snATAC-seq data, the adult mouse forebrain consisted of 52% excitatory neurons, 24% inhibitory neurons, 12% oligodendrocytes and 6% astrocytes and microglia, respectively (Fig. 2d). Since the cell type proportion varies between different forebrain regions, for example cortex and hippocampus^19^, the percentages derived from snATAC-seq represent an average of all forebrain region with numbers in between region-specific values (Extended Data Figure 6d, e; Fig.2e). The predominance of neuronal nuclei derived from adult forebrain tissue was confirmed by flow cytometry analysis using staining against the post-mitotic neuron marker NeuN^18^(Extended Data Fig. 6b). Of note, the proportion of NeuN positive nuclei was lower than the total neuronal proportion derived from snATAC-seq (Extended Data Fig. 6b, e; Fig. 2e).

**Figure 2:**
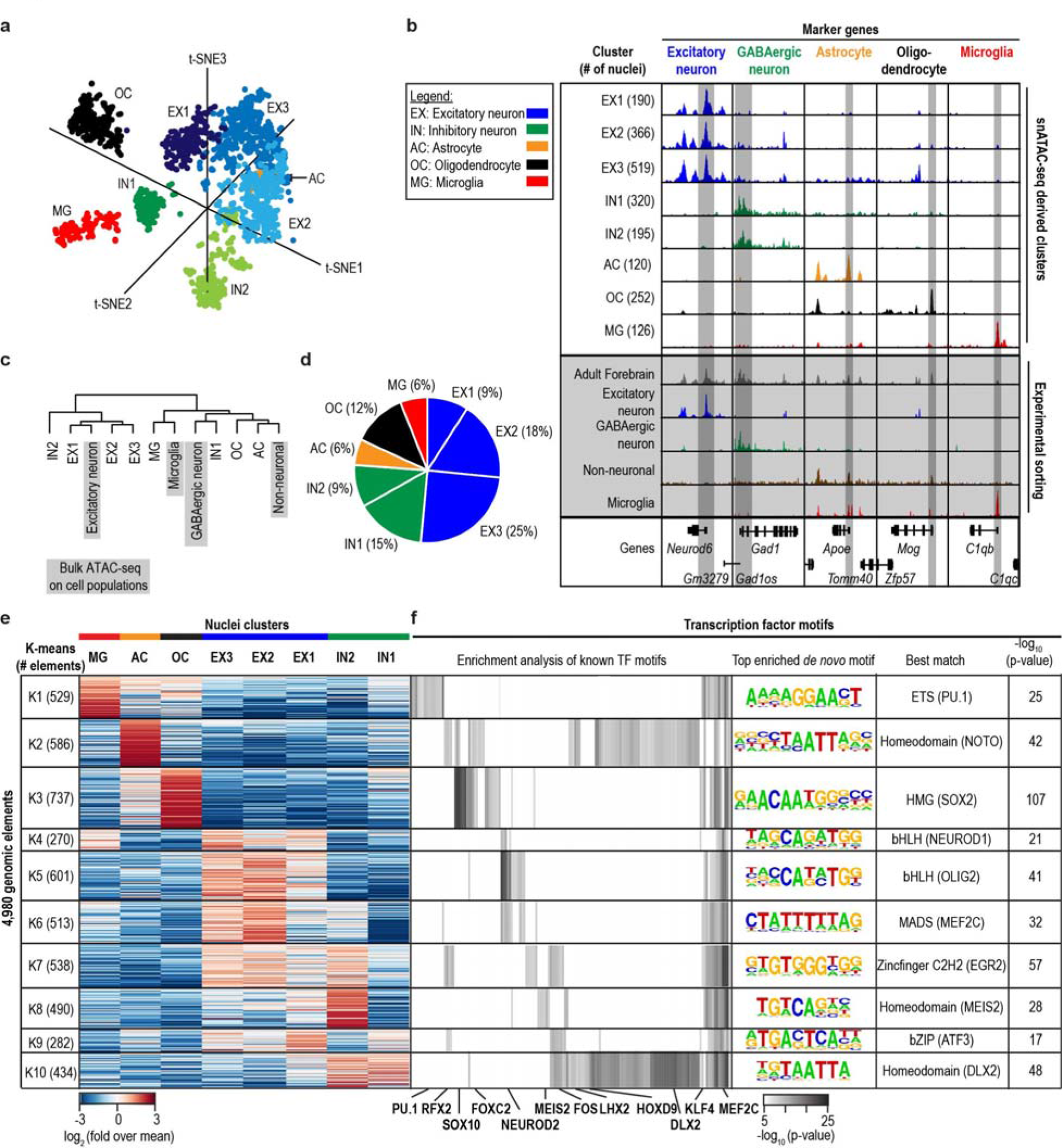
Deconvolution of p56 forebrain and identification of potential master regulators. **a** Clustering of single nuclei of both replicates revealed 8 different cell groups in adult forebrain. **b** Aggregate chromatin accessibility profiles for each cell cluster and bulk ATAC-seq for sorted cell populations or whole forebrain at marker gene loci (Bulk data are grey shaded). **c** Hierarchical clustering of aggregate single cell data and sorted bulk data sets. **d** Cellular composition of adult forebrain derived from snATAC-seq data. **e** K-means clustering of 4,980 genomic elements based on chromatin accessibility and **f** enrichment analysis for transcription factor motifs.

**Extended Data Figure 6:**
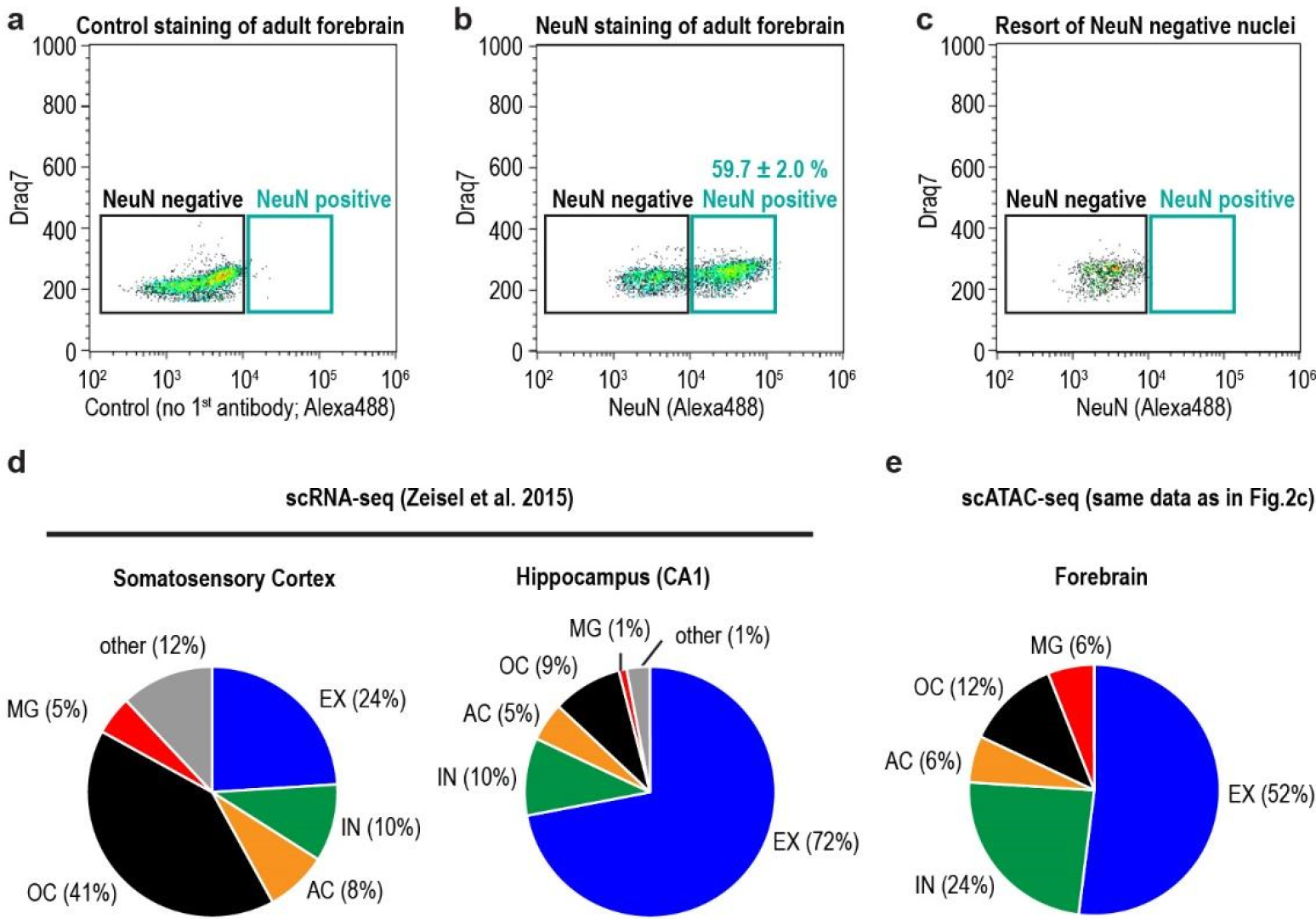
Flow cytometric analysis of adult mouse forebrain and comparison to single cell RNA-seq data from different brain regions. **a-c** Dot blots illustrating nuclei from adult forebrain stained for flow cytometry with Alexa488 conjugated secondary antibodies. **a** Displayed are representative blots for experiments without antigen specific primary antibody and **b** with antibodies recognizing the post-mitotic neuron marker NeuN^18^ (n=3, average ± SEM)**. c** NeuN negative nuclei were sorted for ATAC-seq experiments and purity ( > 98%) was confirmed by flow cytometry of the sorted population. **d** Relative composition of different forebrain regions derived from single cell RNA-seq shows region specific differences^19^**. e** Relative composition derived from snATAC-seq (compare to Fig.2c) of adult forebrain shows values in between.

**Extended Data Figure 7:**
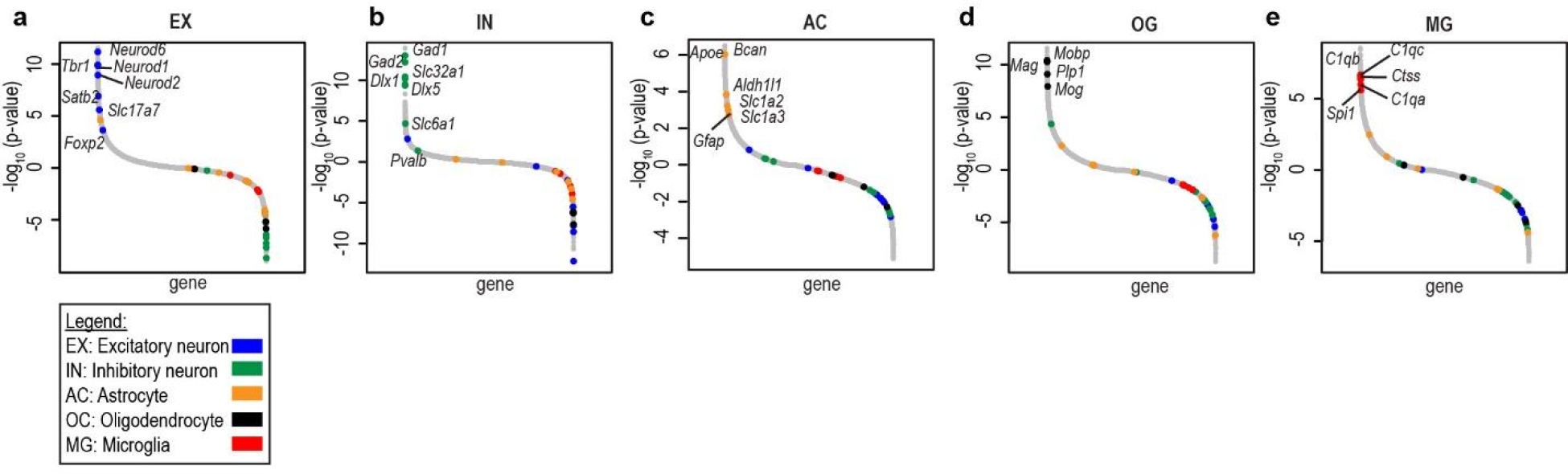
Ranking of gene loci (TSS ± 10kb) compared to other clusters in adult forebrain. Negative binomial test shows enrichment for **a** excitatory neuron markers **b** inhibitory neuron markers **c** astrocyte markers **d** oligodendrocyte markers and **e** microglia markers extending the examples shown in Fig. 2b. Please note for general assignment accessibility profiles for Ex1-3 and IN1/2 were merged, respectively.

To further delineate the *cis*-regulatory landscape in each cell population of the adult forebrain, we plotted the frequency a cis regulatory element was accessible in a nucleus against the cell type specificity index of the element measured by the Shannon entropy of normalized read counts (Extended Data Fig. 8). Overall, proximal promoter elements were accessible in more cell types (Median value of 4.2 *%* for proximal elements vs. 0.4 *%* for distal elements) while the distal enhancer elements showed significantly higher cell type-specificity (Extended Data Fig. 8a, b, d). Next, to identify accessible chromatin regions that distinguish different cell populations, we developed a feature selection method (Methods), and used it to identify a total of 4,980 genomic elements that could separate the 8 nuclei populations in adult mouse forebrain (Fig. 2e, Extended Data Fig. 8c, d). We next performed k-means clustering against the 4,980 genomic regions and conducted motif enrichment analysis of each cluster of elements (Fig. 2e, f, Extended Data Fig. 8d, Supplementary Table 1). As expected, we observed enrichment of known transcription factor motifs in open chromatin in each cell population, including ETS-factor PU.1 for microglia^23^, SOX family of proteins for oligodendrocytes^24^, bHLH factors for excitatory neurons and DLX homeodomain factors for inhibitory neurons (Fig. 2f)^25^. Our analysis also revealed that MEIS binding motif was enriched in a subset of elements specific to IN2. Previous reports showed that MEIS2 plays a major role in generation of medium spiny neurons, the main GABAergic neurons in the striatum^26^. Accordingly, we identified gene loci of *Ppp1r1b* and *Drd1,* markers of medium spiny neurons, to be highly accessible in IN2 but not IN1 (Extended Data Fig. 9)^26^. Next, we asked if we could further separate the excitatory neurons to classes that reflect different anatomical areas. Hierarchical clustering with published bulk ATAC-seq data from different cortical layers and from dentate gyrus^9,27^ showed that clusters EX1-3 might resemble different anatomical regions (Extended Data Fig. 10a). EX1 and EX3 represented upper and lower cortical layers, respectively, whereas EX2 showed properties of dentate gyrus neurons (Extended Data Fig. 10a, b). Regions specific for EX1 and 3 were enriched for motifs from the Forkhead family and EX3 was enriched for motifs recognized by MEF2C (Extended Data Fig. 10c, Supplementary Table 1), which has been shown to play an important role in hippocampus mediated memory^28^.

**Extended Data Figure 8:**
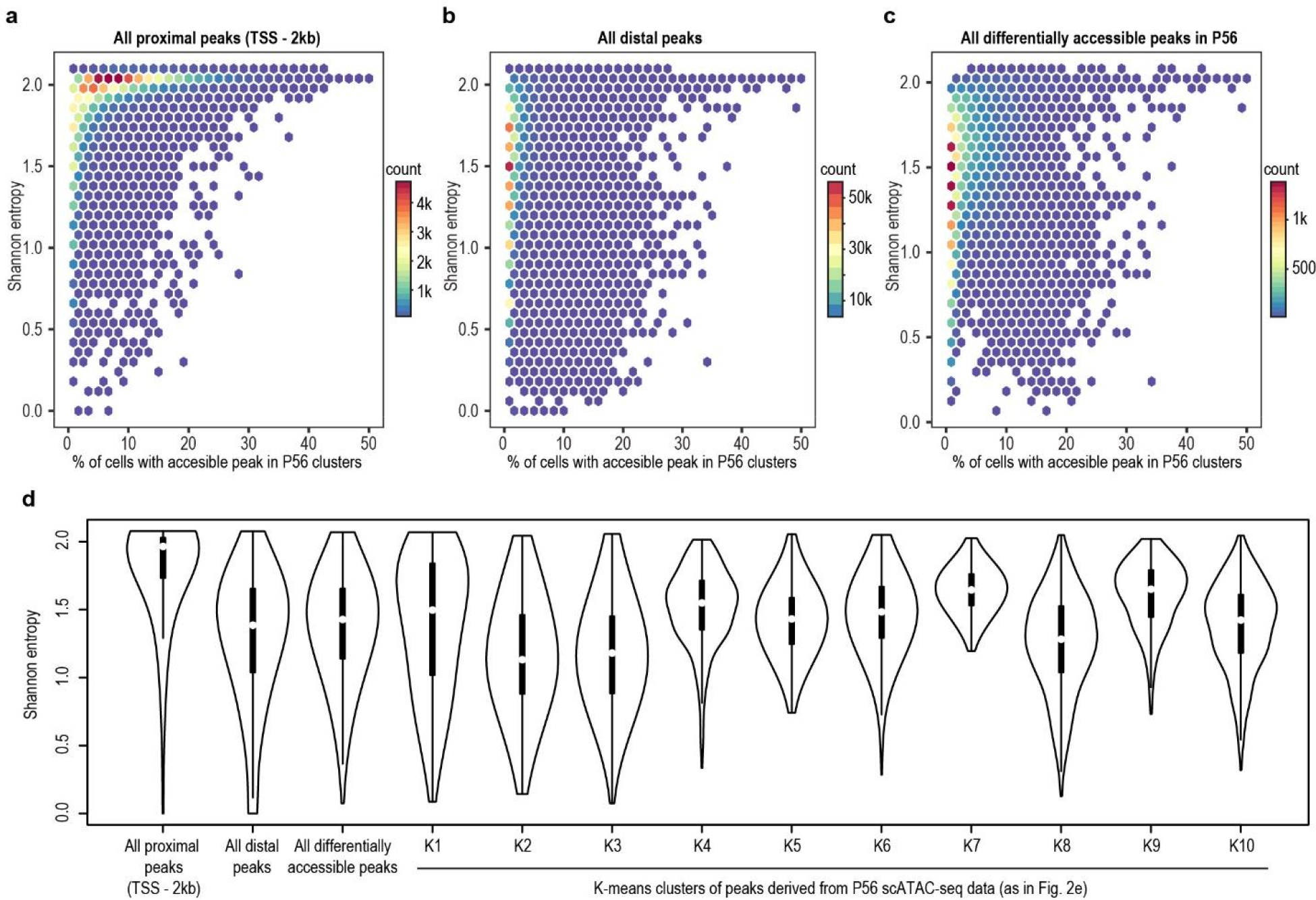
Cell-type specificity of genomic elements and per cell coverage of elements. **a-c** Graphs illustrate cell-type specificity of genomic elements as measured by Shannon entropy based on normalized read counts for each cell-type and percentage of cells in which a genomic element was called accessible as indicated by presence of at least 1 read overlapping with the element a peak. Analysis was performed for the adult forebrain (P56) against **a** TSS-proximal genomic elements (TSS-2kb), **b** distal elements and **c** the subset of genomic elements that separated two cell clusters. **d** Violin plots illustrate higher cell-type specificity for distal elements compared to proximal elements indicated by significantly lower Shannon entropy value (p < 2.2e-16). In addition, distribution of all genomic elements that separate two clusters as well as subsets representing subsets identified from k-means clustering of genomic elements depending on chromatin accessibility in adult forebrain (related to Fig. 2e). TSS: transcriptional start site.

**Extended Data Figure 9:**
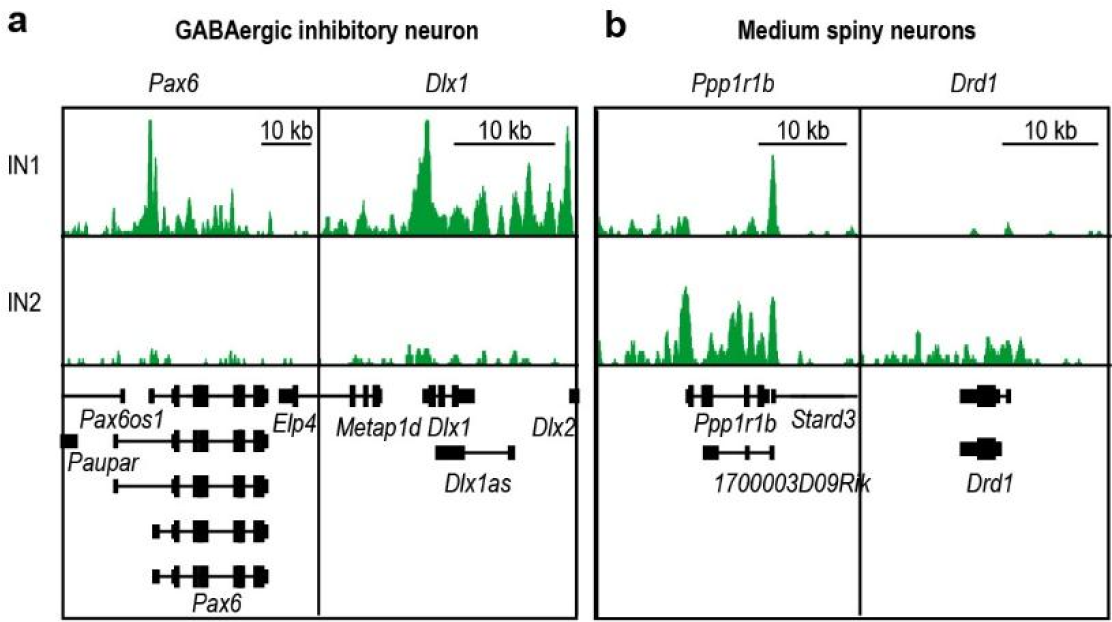
Distinct chromatin accessibility profiles of two GABAergic neuron clusters. IN2 is depleted for *Pax6* and *Dlx1* (**a**) but enriched for markers of medium spiny neurons as compared to IN1 cluster (**b**).

**Extended Data Figure 10:**
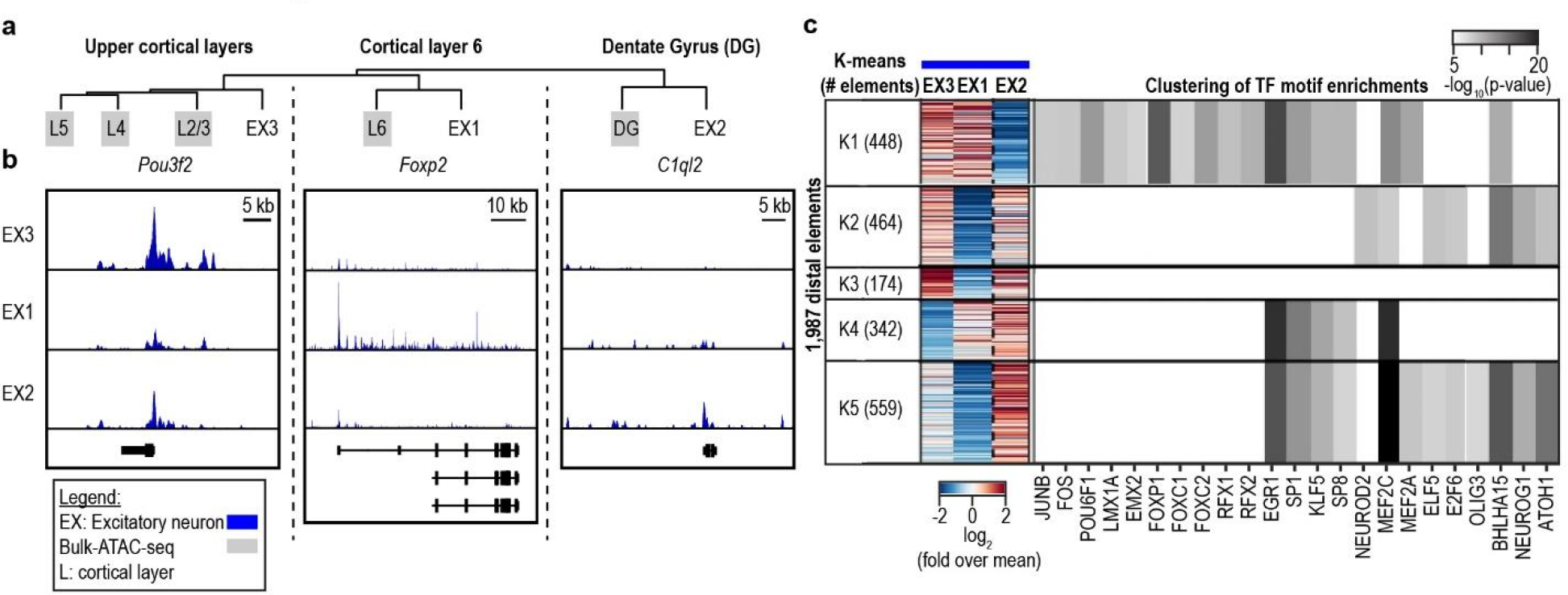
Sub-classification of excitatory neurons into hippocampal and cortical neuron types. **a** Hierarchical clustering of aggregate single cell data for excitatory neuron cluster and sorted bulk data sets corresponding to different anatomical regions (grey shaded). **b** Chromatin accessibility at marker gene loci. **c** K-means clustering of promoter distal genomic elements and enrichment analysis for transcription factor motifs.

We next examined the snATAC-seq profiles derived from foetal mouse forebrains from seven developmental stages (Fig. 1b) that include key events from the onset of neurogenesis to gliogenesis^29^. From 12,733 high-quality single nuclei ATAC-seq profiles we identified 12 distinct sub-populations (Fig. 3a) that exhibit dynamic abundance through development (Fig. 3a-c). Based on accessibility profiles at gene loci of known marker genes, we assigned these cell populations to radial glia, excitatory neurons, inhibitory neurons, astrocytes and erythromyeloid progenitors (EMP) (Fig. 3b)^23,30^. Interestingly, the EMP cluster was restricted to E11.5, whereas the astrocyte cluster was present after E16.5 and expanded dramatically around birth (Fig. 3b, c)^23,29^, highlighting two developmental processes: invasion of myeloid cells into the brain prior to neurogenesis, and gliogenesis succeeding neurogenesis after E16.5^29^. Mature excitatory neurons (eEX2) were indicated by increased accessibility at the post-mitotic neuron marker gene *Neurod6* and absence of signal at the Notch effector *Hes5,* a marker gene for neuronal progenitors (Fig. 3b, c)^29,30^. This cell type expanded in abundance between E12.5 and E13.5 and followed the emergence of early differentiating neurons (eEX1, Fig. 3b, c). Remarkably, inhibitory-neuron-like cells were already present at E11.5 (Fig. 3b).

**Figure 3:**
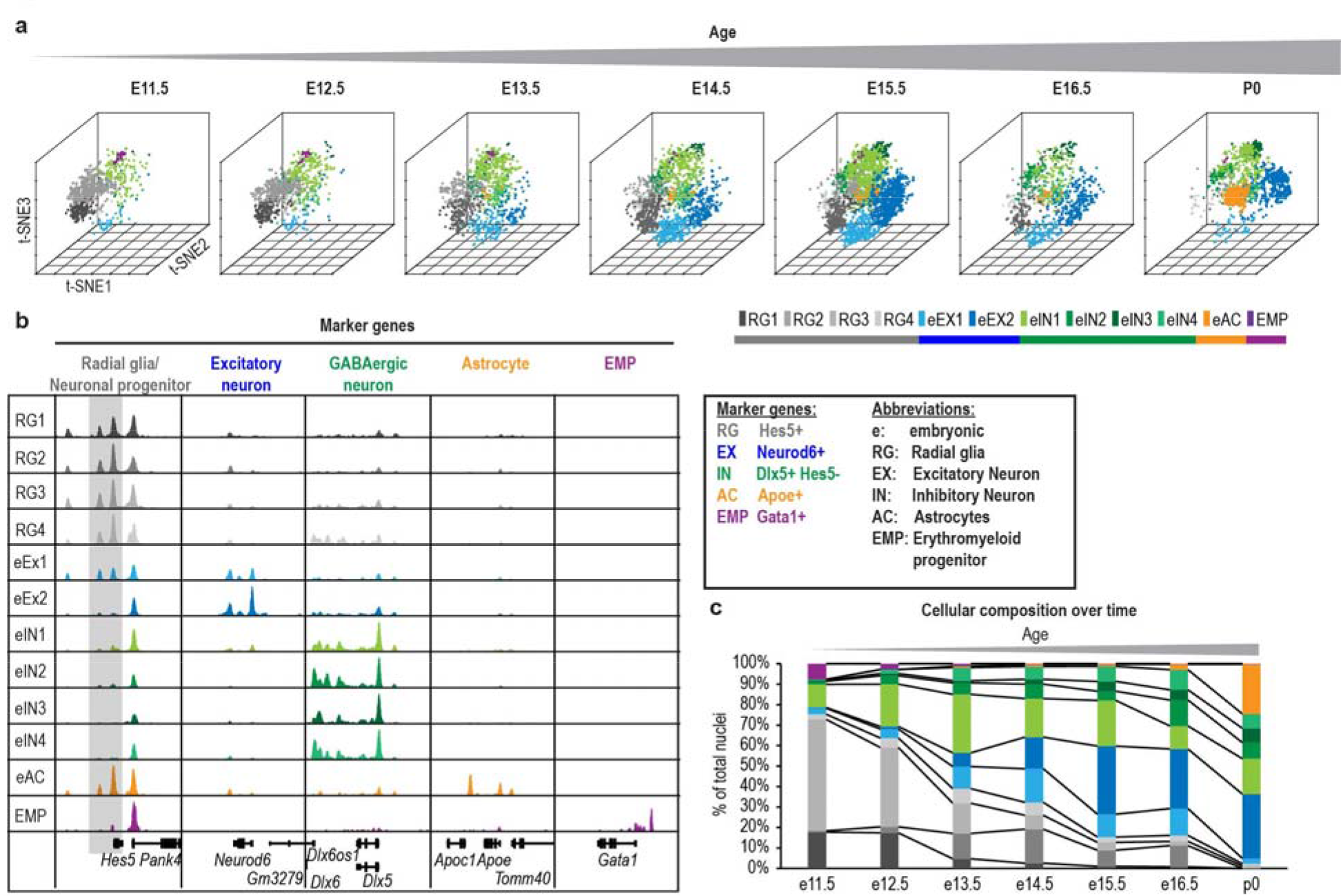
SnATAC-seq analysis reveals cellular heterogeneity during embryonic forebrain development. **a** Clustering of single nuclei from both replicates revealed 12 different cell groups with changing relative abundance. **b** Aggregate chromatin accessibility profiles for cell clusters and at marker gene loci used to assign cell types. For better visualization, *Hes5* gene locus is grey shaded. **c** Quantification of cellular composition during forebrain development.

To identify the transcriptional regulatory sequences in each sub-population, we identified 16,364 genomic elements that show cell-population-specific chromatin accessibility and can best separate the sub-cell populations (Fig. 4a, Supplementary Table 2). We clustered these elements using k-means and performed gene ontology analysis of each cluster using the GREAT^31^. We also conducted *de novo* motif search for each group of elements to uncover transcription factor motifs enriched in cell type-specific open chromatin regions (Fig. 4b, c). Our analysis showed that genomic elements that were mostly associated with radial glia like cell groups (Fig. 4a, RG1-4) fell into regulatory regions of genes involved in early forebrain developmental processes including “Forebrain regionalization” (Fig. 4b, K1), “Central nervous system development” (Fig. 4b, K3) or “Forebrain development” (Fig. 4b, K5). These elements were enriched for homeobox motifs corresponding to LHX-transcription factors including LHX2 (Fig. 4c, K1, 3, 5), which is critical for generating the correct neuron number by regulating proliferation of neural progenitors^32^ and for temporal promoting of neurogenesis over astrogliagenesis^33^. Remarkably, one of these cluster was also enriched for both the proneural bHLH transcription factor ASCL1 (*Mash1*) and its co-regulator POU3F3 *(Brn1)* (Fig. 4c, K5)^34^. ASCL1 has been described to be required for normal proliferation of neural progenitor cells^35^ and implicated in a DLX1/2 associated network that promotes GABAergic neurogenesis^36^. In line with this, associated genomic elements were also accessible in one inhibitory neuron cluster (eIN2, Fig. 4c, K5).

**Figure 4:**
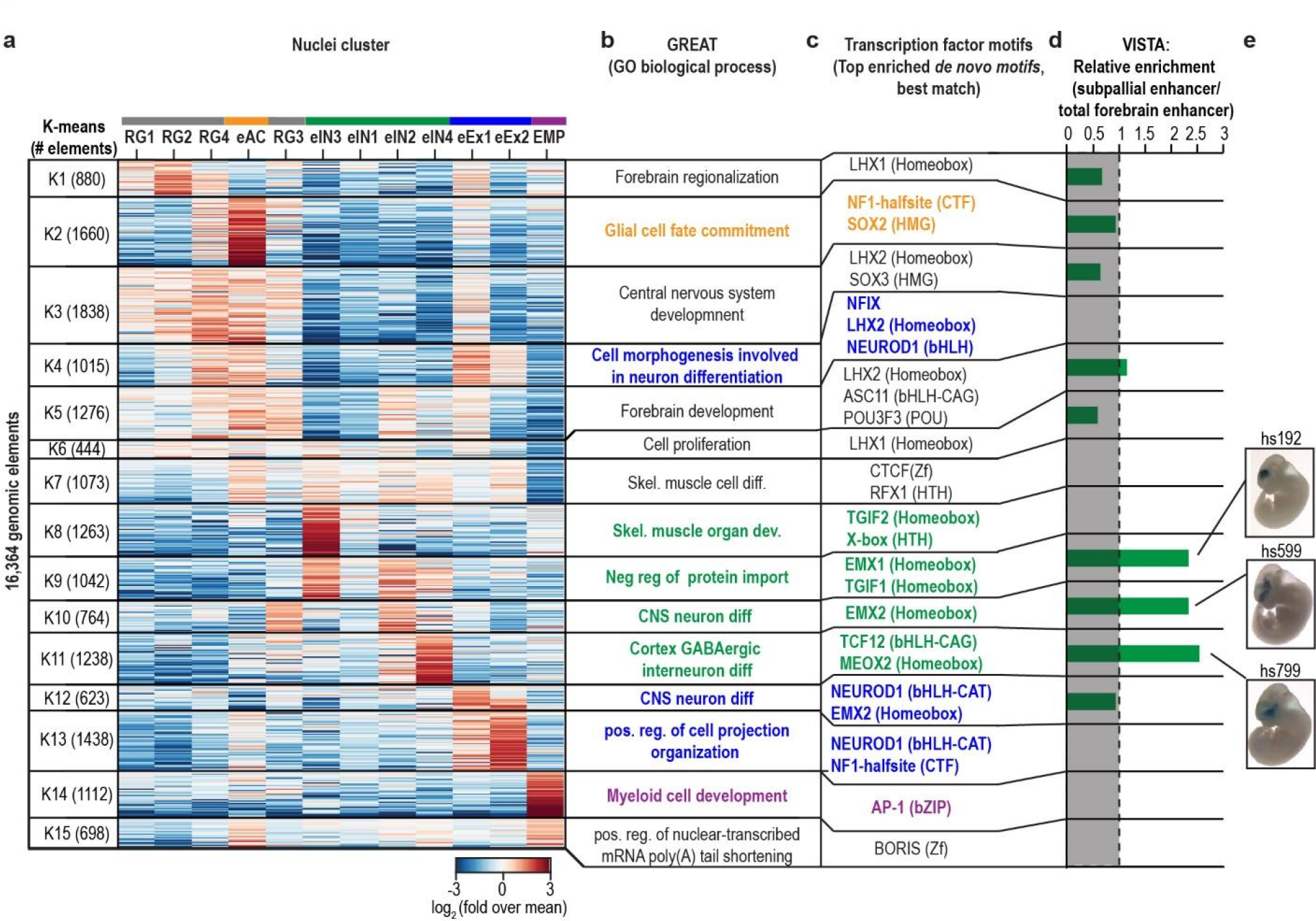
SnATAC-seq revealed genomic elements and transcriptional regulators of lineage specification in the developing forebrain. **a** K-means clustering of 16,364 genomic elements based on chromatin accessibility. **b** Gene ontology analysis using GREAT and **c** transcription factor enrichment. **d** Enrichment of enhancers that were functionally validated as part of the VISTA database. **e** Representative pictures of transgenic mouse embryos showing LacZ reporter gene expression under control of the indicated subpallial enhancers. Pictures were downloaded from the VISTA database^44^.

We identified transcriptional regulators that were specifically associated either with neurogenesis or gliogenesis during forebrain development. For example, the early astrocyte (eAC)-specific elements were located in open chromatin regions near genes involved in “glia cell fate commitment” and the top enriched transcription factor motif was NF1-halfsite (Fig.4a-c, K2). Previous studies showed that NF1 transcription factor NF1A alone is capable for specifying glia cells to the astrocyte lineage^24^. NFIX is another NF1 family member with proneural function^37^. This motif is enriched together with the bHLH transcription factor NEUROD1 binding sites mainly in open chromatin regions found in the excitatory neuron cell population (Fig4c, K4, 12, 13)^30^. Based on chromatin accessibility profiles at marker gene loci, we have previously assigned two cell clusters to excitatory neuron lineage (eEX1, eEX2, Fig.3b). Compared to cluster eEX2, eEX1 showed increased accessibility at both radial glia associated open chromatin (Fig.4a, K4; Fig.3b) and chromatin regions associated with “CNS neuron differentiation” (Fig.4a, K12). In addition, eEX1 nuclei preceded the emergence of eEX2 nuclei during development (Fig.3c). These findings indicate that eEX1 might represent a transitional state during excitatory neuron differentiation.

The bHLH transcription factor family consists of several subfamilies that recognize different DNA motifs^38^. NEUROD1 is part of a subfamily that binds to a central CAT motif whereas other factors such as TCF12 preferentially bind to a CAG motif^38^. Our snATAC-seq data revealed an enrichment of the TCF12-binding motif in regions associated with “Cortex GABAergic interneuron differentiation” in contrast to the excitatory neuron associated enrichment for NEUROD1 (Fig.4a-c, K4, 11, 12, 13)^25,30,39^.

Analysis of inhibitory neuron cluster eIN3 specific genomic elements showed a remarkable enrichment for genes associated with “Skeletal muscle organ development” (Fig4a, b, K8). More detailed analysis revealed that the underlying genes were transcriptional regulators *Mef2c/d* and *Foxp1/2* as well as the dopamine receptors Drd2/3 indicating differentiating striatal medium spiny neurons^40,41^. This finding was consistent with the enrichment for MEIS-homeodomain factors in these regions (Fig.4c, K8) comparable to the medium spiny neuron cluster in adult forebrain (Fig.2e, f, K8; Extended Data Fig.9).

Lastly, genomic elements specific to the EMP cluster were associated with genes involved in “Myeloid cell development” (Fig.4a-c, K14) and enriched for motifs of the ubiquitous AP-1 transcription factor complexes that have been described to play a role in shaping the enhancer landscape of macrophages^42^.

Next, we attempted to identify dynamic elements within a given cell clusters (Extended Data Fig.11). Our analysis revealed between 41 and 2,114 dynamic genomic elements for each cell type (Extended Data Fig.11c-g). Regions that are more accessible after birth (P0) compared to early time points were enriched for the RFX1 motif in the GABAergic neuron including the cluster eIN1 as well as in the excitatory neuron cluster eEX2 (Extended Data Fig.11d, e) indicating a general role of the evolutionary conserved RFX factors in perinatal adaptation of brain cells. Several family members including RFX1 are expressed in the brain and have been implicated to regulate cilia e.g. in sensory neurons^43^.

**Extended Data Figure 11:**
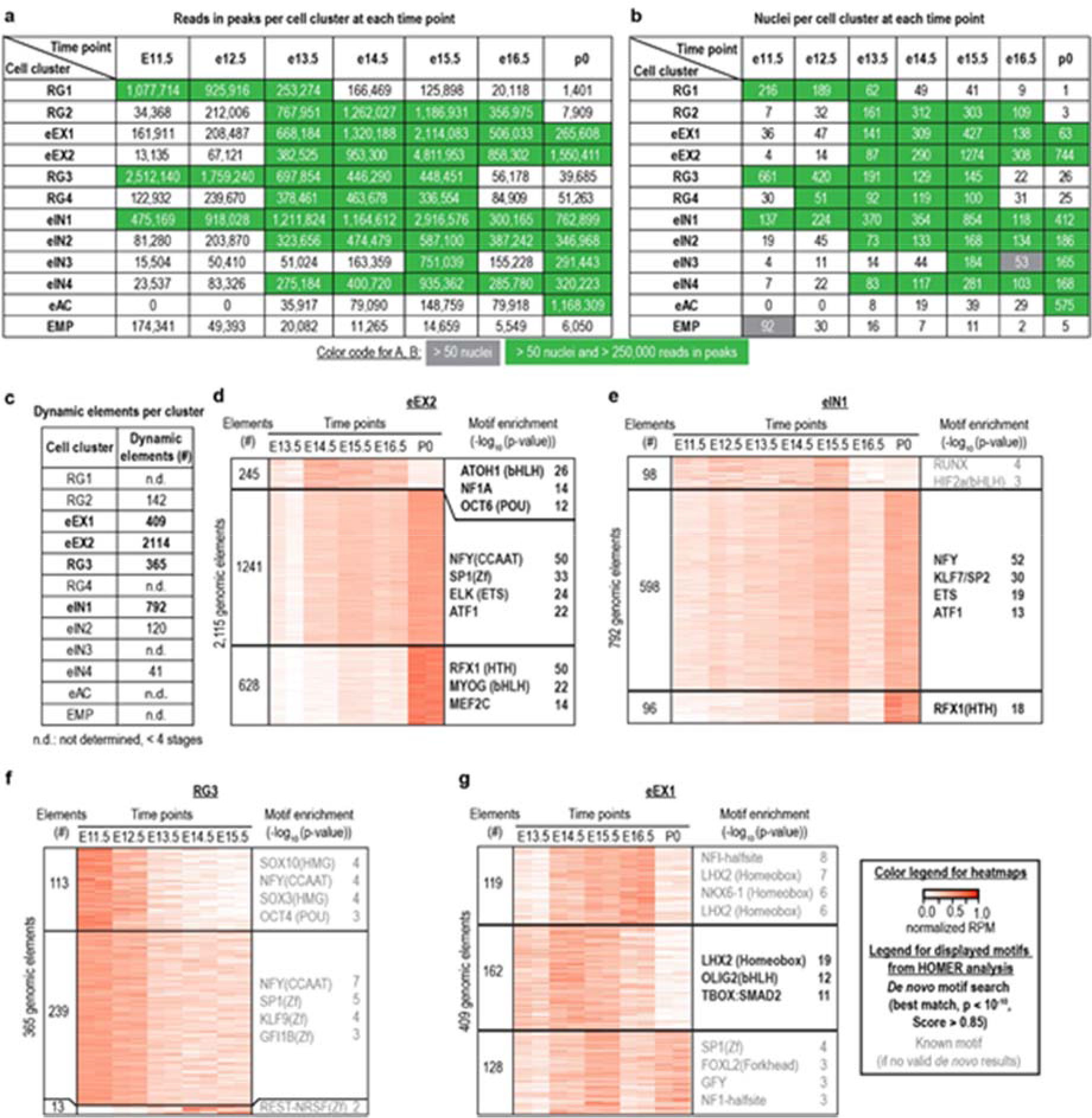
Dynamics of chromatin accessibility within distinct cell groups. **a** Number of reads in peaks per developmental time point for a specific nuclei cluster. **b** Number of nuclei per time point for a specific nuclei cluster. For analysis of dynamics only cell clusters with > 3 stages with > 50 nuclei and > 250,000 reads in peaks were considered. **c** Overview of dynamic elements identified per cell cluster (see methods) **d-g** K-means clustering and motif enrichment analysis for nuclei clusters with > 200 dynamic genomic elements.

While assessment of open chromatin plays an important role in predicting regulatory elements in the genome^1,3^, it does not provide direct information of functional activity. To address this point, we asked if cell-type specific transposase accessible chromatin in the embryonic forebrain overlaps with known enhancers validated by transgenic mouse assays^44^. We focused our analysis on all genomic elements with validated functional activity in the forebrain and a subset shown to be active only in the subpallium^45,46^ The subpallium is a brain region that gives rise to GABAergic and cholinergic neurons^45^. In total, 63.1 *%* (275/436) of all forebrain enhancer and 64.8% (59/91) of subpallial enhancer were represented in our subset of genomic elements, respectively, indicating a high degree of sensitivity. Next, we calculated the relative enrichment of subpallial enhancers over total forebrain enhancers for each cluster. Remarkably, subpallial enhancers were only enriched in clusters K9-11, which were assigned to the GABAergic neuron lineage (Fig.4d, e). This analysis confirms a high specificity and sensitivity of snATAC-seq experiments in identifying sub-cell populations and their underlying regulatory elements.

Taken together, we demonstrate here that snATAC-seq can be used to dissect heterogeneity and delineate gene regulatory sequences in complex tissues such as forebrain. Using this strategy, we were able to resolve the heterogeneity of primary tissue samples, and uncover both the cell types and the regulatory elements in each cell type without prior knowledge. The snATAC-seq approach will be a valuable tool for analysis of tissue biopsies and will help to pave the way to a better understanding of gene regulation in mammals.

## AUTHOR CONTRIBUTIONS

Study was conceived and designed by B.R., S.P., R.F.; Study was overseen by B.R.; Experiments performed by S.P., B.C.S, H.H.; Tissue collection by V.A., D.E.D., S.P.; Sequencing performed by S.K.; Computational strategy developed by R.F. Data analysis performed by S.P., R.F., R.R., Y.Z., D.U.G; Manuscript written by S.P., R.F. and B.R. with input from all authors.

## ACKNOWLEDGMENTS

This study was funded in part by the National Human Genome Research Institute as part of the Encyclopedia of DNA Elements (ENCODE) project (U54HG006997) and supported by funding from the Ludwig Institute for Cancer Research and NIH (2P50 GM085764). S.P. was supported by a postdoctoral fellowship from the Deutsche Forschungsgemeinschaft (DFG, PR 1668/1-1). RR was supported by a Ruth L. Kirschstein National Research Service Award NIH/NCI T32 CA009523. We thank Bin Li for bioinformatic support. We thank Molly He and Trina Osothprarop for providing the Tn5 enzyme. We thank Derek Gao for sequencing on the MiSeq. This study was supported by the NIH grant U01MH098977 to K.Z. Research conducted at the E.O. Lawrence Berkeley National Laboratory was performed under U.S. Department of Energy Contract DE-AC02-05CH11231, University of California.

## METHODS

### Mouse tissues

All animal experiments were approved by the Lawrence Berkeley National Laboratory Animal Welfare and Research Committee or the University of California, San Diego, Institutional Animal Care and Use Committee. Forebrains from embryonic mice (E11.5-E16.5) and early postnatal mice (P0) were dissected from one pregnant female or one litter at a time and combined. For breeding, animals were purchased from Charles River Laboratories (C57BL/6NCrl strain) or Taconic Biosciences (C57BL/6NTac strain) for E14.5 and P0. Breeding animals for other time points were received from Charles River Laboratories (C57BL/6NCrl). Dissected tissues were flash frozen in a dry ice ethanol bath. For the adult time point (P56), the forebrain from 8-week old male C57BL/6NCrl mice (Charles River Laboratories) were dissected and flash frozen in liquid nitrogen separately. Tissues were pulverized in liquid nitrogen using pestle and mortar. For each time point two biological replicates were processed (n = 2 per time point).

### Transposome generation

To generate A/B transposomes, A and B oligos were annealed to common pMENTs oligos (95°C 2 min, 14°C ∞ (cooling rate: 0.1°C/s)) separately (Supplementary Table 2). Next, barcoded transposons were mixed in a 1:1 molar ratio with unloaded transposase Tn5 which was generated at Illumina. Mixture was incubated for 30 min at room temperature. Finally, A and B transposomes were mixed. For combinatorial barcoding we used 8 different A transposons and 12 distinct B transposons which eventually resulted in 96 barcode combinations (Supplementary Table 2)^49^.

### Combinatorial barcoding assisted single nuclei ATAC-seq

Combinatorial ATAC-seq was performed as described previously with modifications^5^. 5-10 mg frozen tissue was transferred to a 1.5 ml Lobind tube (Eppendorf) in 1 ml NPB (5 % BSA (Sigma), 0.2 % IGEPAL-CA630 (Sigma), cOmplete (Roche), 1 mM DTT in PBS) and incubated for 15 min at 4 °C. Nuclei suspension was filtered over a 30 µm Cell-Tric (Sysmex) and centrifuged for 5 min with 500 x g. Nuclei pellet was resuspended in 500 µl of 1.1x DMF buffer (36.3 mM Tris-acetate (pH = 7.8), 72.6 mM K-acetate, 11 mM Mgacetate, 17.6 % DMF) and nuclei were counted using a hemocytometer. Concentration was adjusted to 500 /µl and 4500 nuclei were dispensed into each well of a 96 well plate. For tagmentation, 1 µl barcoded Tn5 transposome (0.25 µM, Supplementary Table 2)^49^ was added to each well, mixed 5 times and incubated for 60 min at 37°C with shaking (500 rpm). To quench the reaction 10 µl 40 mM EDTA were added to each well and plate was incubated at 37°C for 15 min with shaking (500 rpm). 20 µl sort buffer (2 % BSA, 2 mM EDTA in PBS) were added to each well and all wells combined afterwards. Nuclei suspension was filtered using a 30 µm CellTric (Sysmex) into a FACS tube and 3 µM Draq7 (Cell Signalling) was added. Using a SH800 sorter (Sony) 25 nuclei were sorted per well into 4 96-well plates (total of 384 wells) containing 18.5 µl EB (50 pM Primer i7 (Supplementary Table 2), 200 ng BSA (Sigma)). Sort plates were shortly spun down. After addition of 2 µl 0.2 % BSA, samples were incubated at 55°C for 7 min with shaking (500 rpm). 2.5 µl 10% Triton-X was added to each well to quench SDS. Finally, 2 µl 25 µM Primer i5 (Supplementary Table 2) and 25 µl NEBNext® High-Fidelity 2X PCR Master Mix (NEB) and samples were PCR amplified for 11 cycles (72°C 5 min, 98°C 30 s,[ 98°C 10 s, 63°C 30 s, 72°C 60 s] x 11, 72°C ∞). Following PCR, all wells were combined (around 15.5 mL) and mixed with 80 ml PB including pH-indicator (1:2500, Qiagen) and 4 ml Na-Acetate (3 M, pH = 5.2). Purification was carried out on 4 columns following the MinElute® PCR Purification Kit manual (Qiagen). DNA was eluted with 15 µl EB and eluate from all four columns was combined in a LoBind Tube (Eppendorf). For Ampure XP Bead (Beckmann Coulter) cleanup 170 µl EB buffer and 110 µl Ampure XP Beads (0.55x) were added to 30 µl eluate. After incubation at room temperature for 5 min and magnetic separation supernatant was transferred to a new tube and another 190 µl Ampure XP Beads (1.5x) were added. After incubation beads were washed twice on the magnet using 500 µl 80 % EtOH. After drying the beads for 7 min at room temperature library was eluted with 20 µl EB (Qiagen). Libraries were quantified using Qubit fluoromoeter (Life technologies) and nucleosomal pattern was verified using Tapestation (High Sensitivity D1000, Agilent). 25 pM library was loaded per lane of a HiSeq2500 sequencer (Illumina) using custom sequencing primers (Supplementary Table 2)^49^ and following read lengths: 50 + 43 + 37 + 50 (Read1 + Index1 + Index2 + Read2). The first 8 bp of Index1 correspond to the p7 barcode and the last 8 bp to the i7 barcode. The first 8 bp of Index2 correspond to the i5 barcode and the last 8 bp to the p5 barcode. Since Index1 and 2 each contain 2 barcodes separated by a common linker sequence, we generated a spike-in library using different transposon and PCR primer sequences to balance the bases within each detection cycle (Supplementary Table 2). For the human-mouse mixture experiment, E15.5 forebrain and GM12878 nuclei were mixed in a 1:1 ratio prior to tagmentation. Samples were processed as above with the exceptions that just 96 wells were used after nuclei sorting and PCR amplification was performed for 13 cycles. The final library was loaded at 15 pM and sequenced using a MiSeq (Illumina) with following read lengths: PE 44 + 43 + 37 +44 (Read1 + Index1 + Index2 + Read2).

### Cell culture

GM12878 (Coriell Institute for Medical Research) cells were cultured in RPMI1640 medium (Thermo Fisher Scientific) containing 2 mM L-glutamine (Thermo Fisher Scientific), 15% foetal bovine serum (Gemini Bioproducts) and 1 % Penicillin-Streptomicin (Thermo Fisher Scientific) in T25 Flasks (Corning) at 37°C under 5% carbon dioxide. For the snATAC-seq mixture experiment, cells were harvested by centrifugation, washed with PBS (Thermo Fisher Scientific) and resuspended in NPB (5 % BSA (Sigma), 0.2 % IGEPAL-CA630 (Sigma), cOmplete (Roche), 1 mM DTT in PBS). Samples were incubated 5 min at 4 °C and finally nuclei were pelleted by centrifugation (500g, 5min, 4 °C). Nuclei pellet was resuspended in 500 µl of 1.1x DMF buffer (36.3 mM Tris-acetate (pH = 7.8), 72.6 mM K-acetate, 11 mM Mg-acetate, 17.6 % DMF) and nuclei were counted using a hemocytometer.

### NeuN negative sorting

10 mg adult forebrain tissue (P56) were resuspend in 500 µl lysis buffer (0.5% BSA, 0.1% Triton-X, cOmplete (Roche), 1 mM DTT in PBS) and incubated for 10 min at 4°C. After spinning down (5 min, 500 x g) sample was resuspended in 500 µl staining buffer (0.5% BSA in PBS). Nuclei suspension was incubated with anti-NeuN antibody (1:5000, MAB377, Lot 2806074, EMD Millipore) for 30 min at 4°C. After centrifugation nuclei were resuspend in 500 µl staining buffer (0.5% BSA in PBS) containing anti-mouse Alexa488-antibody (1:1000, A11001, Lot 1696425, Thermo Fisher Scientific). After incubate for 30 min at 4°C, nuclei were pelleted (5 min 500 x g) and resupended in 700 ul sort buffer (1% BSA, 1mM EDTA in PBS). After filtration into a FACS tube 5 ul DRAQ7 (Cell Signalling Technologies) was added and NeuN-negative nuclei were sorted using a SH800 sorter (Sony) into 5% BSA (Sigma) in PBS.

### ATAC-seq

ATAC-seq was performed on 20,000 sorted nuclei as described previously with minor modifications^50^. After adding IGEPAL-CA630 (Sigma) in a final concentration of 0.1 % nuclei were pelleted for 15 min at 1000 x g. Pellet was resupended in 19 µl 1.1x DMF buffer (36.3 mM Tris-acetate (pH = 7.8), 72.6 mM K-acetate, 11 mM Mg-acetate, 17.6 % DMF). After addition of 1 µl Tn5 transposomes (0.5 µM) tagmentation was performed at 37°C for 60 min with shaking (500 rpm). Next, samples were purified using MinElute columns (Qiagen), PCR-amplified for 8-10 cycles with NEBNext® High-Fidelity 2X PCR Master Mix (NEB, 72°C 5 min, 98°C 30 s,[ 98°C 10 s, 63°C 30 s, 72°C 60 s] x cycles, 72°C ∞). Amplified libraries were purified using MinElute columns (Qiagen) and Ampure XP Bead (Beckmann Coulter). Sequencing was carried out on a HiSeq2500 or 4000 (50 bp PE, Illumina).

## DATA ANALYSIS

### Single nuclei ATAC-seq data processing pipeline

1. <u>Alignment:</u> Paired-end sequencing reads were aligned to mm10 reference genome using Bowtie2 in paired-end mode with following parameters “bowtie2-p 5-t-X2000 ‒‒no-mixed ‒‒no-discordant”^51^.
2. <u>Alignment filtration:</u> Non-uniquely mapped (MAPQ < 30) and improperly paired (flag = 1804) alignments were filtered.
3. <u>Barcode error correction:</u> Each full barcode consists of four separate 8 bp long indexes (i5, i7, p5, p7). Reads with barcode combinations containing more than 1 mismatch for any index were removed. Barcodes with ≤ 1 mismatch were changed to its closest barcode.
4. <u>Reads separation:</u> Reads were separated into individual cells based on the barcode combination (Extended Data Table 1, Supplementary Table 2).
5. <u>Mark and remove PCR duplicates:</u> For individual cells, we sorted reads based on the genomic coordinates using “samtools sort”^52^, then marked and removed PCR duplicates using Picard tools (MarkDuplicates, https://broadinstitute.github.io/picard/).
6. <u>Mitochondrial reads removal:</u> Reads mapped to the mitochondrial genome were filtered.
7. <u>Adjusting position of Tn5 insertion:</u> All reads aligning to the + strand were offset by +4 bp, and all reads aligning to the-strand were offset-5 bp^53^.
8. <u>Quality assessment of each single cell:</u> Calculate coverage of constitutively accessible promoters (promoters that are accessible across all tissues/cell line from ENCODE DHS), number of reads and signal-over-noise ratio estimated by “reads in peaks” ratio for each cell.
9. <u>Cell selection:</u> We only kept cells that pass our threshold (1) coverage of constitutively accessible promoter > 10%; 2) number of reads > 1,000; 3) reads in peak ratio greater than estimation from corresponding bulk ATAC-seq level (Zhao et al. manuscript in preparation).
10. <u>Replicates separation:</u> Selected cells were separated into two replicates based on the predefined barcode combination.

### Single nuclei ATAC-seq cluster analysis

Cluster analysis partitions cells into groups such that cells from the same group have higher similarity than cells from different groups. Here, we developed a pipeline to obtain cell clusters. We first generated a catalogue of accessible chromatin regions using bulk ATAC-seq data (Zhao et al. manuscript in preparation) and created a binary accessible matrix. Chromatin sites were 1 for a given cell if there was a read detected within the peak region. Next, we calculated pair-wise Jaccard index between every two cells on the basis of overlapping open chromatin regions. Next, we applied a non-linear dimensionality reduction method (t-SNE) to map the high-dimensional structure to a 3-D space^14^. This transforms high-dimensional structures to dense data clouds in a low-dimensional space, allowing partitioning of cells using a density-based clustering method^15^. We then identified the optimal number of cell clusters using the Dunn index^54^. Finally, we compared our cluster results to those of “shuffled” to further verify our cluster result is not driven by library complexity or other confounding factors.

1. <u>Determining accessible chromatin sites in single cells</u> To catalogue accessible chromatin sites in individual cells, we first created a reference map of open chromatin sites determined by bulk ATAC-seq (Zhao et al. manuscript in preparation). The chromatin accessibility maps from different time points (from E11.5 to P56) were merged into a single reference file using BEDtools^55^. For clustering of single cells, we have tested clustering performance using accessible promoters (2kb upstream of TSS) and distal elements, respectively, and found that clusters by distal elements outperformed promoters with lower Kullback-Leibler divergence (Extended Fig. 5). Therefore, we decided to only focus on distal genomic elements as features to perform clustering. Reads in individual cells overlapping with accessible sites were identified. We generated an accessible matrix of the reads counts overlapping each individual accessible sites (columns) in each cell (row).
2. <u>Binary Accessible Matrix</u> We next converted the chromatin accessibility matrix to a binary matrix *M_N×D_* in which *M_ij_* is 1 if any read in cell *i* mapped to region *j*.
3. <u>Jaccard Index Matrix</u> Jaccard index matrix *J_N×N_* were calculated between every two cells in which *J_ij_* measures the commonly shared open chromatin regions between cell *C_i_* and *C_j_* as following:

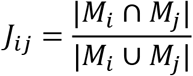

Diagonal elements of *J_N×N_* are set to be 0 as required by t-SNE analysis.
4. <u>Dimensionality reduction using t-SNE</u> Using Jaccard index matrix *J_N×N_* as input, we next applied t-SNE to map the N-dimensional data to a 3-D space^14^. Since t-SNE has a non-convex objective function, it is possible that different runs yield different solutions^14^. Thus, we ran t-SNE several times with different initiations and used the result with the lowest Kullback-Leibler divergence and best visualization. In a previous study sequencing depth was a confounding factor and highly correlated with the first principle component of PCA analysis (Pearson correlation >0.95)^5^. However, we did not observe correlation between sequencing depth and any of the t-SNE dimension. We expected that the coherent structure of the open chromatin landscape of cells with high similarity would rely on a continuous and smooth 3-D structure and cells for different groups would locate to distinct parts of the plot. We used t-SNE to transform the high-dimensional structures to dense data clouds in the 3-D space^14^. Finally, we applied a density-based clustering method to identify different cell populations within the embedded 3-D space^15^.
5. <u>Density-based clustering</u> We applied a density-based clustering method to partition cells into groups in the embedded 3-D space^15^. The method identifies cluster centres that are characterized by two properties: 1) high local density *ρ_i_* and 2) large distance *δ_i_* from points of higher density, which are centers of the clusters^15^. Any cells that showed values above defined thresholds (*ρ_0_,δ_0_*) were considered as centers of cluster. Next, the rest of cells were assigned to the center as described here^15^. Clearly, different thresholds (*ρ_0_,δ_0_*) will generate different number of clusters. To find the optimal number of clusters, we adopted the method developed by Habib et al to evaluate the quality of different cluster results^54^.
6. <u>Number of clusters</u> In detail, Habib’s method applied the Dunn index (Algorithm 2) to quantify the quality of cluster result as following^48^:

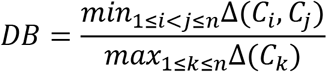

in which Δ(*C_i_, C_j_*) represents the inter-cluster distance between cluster *C_i_* and *C_j_*, Δ(*C_k_*) represents the intra-cluster distance of cluster *C_k_*. We used the “MaxStep” distance (Algorithm 1) also developed by Habib et al to calculate the distance for Dunn index^54^. Finally, we iterated all possible (*ρ_0_,δ_0_*) combinations that yield different clusters and calculated its Dunn index. The clustering result with the highest Dunn index was chosen as final cluster (Algorithm 3).

~~~
**Algorithm 1:** calculate max step distance (MaxStep) (adopted from *Habib* et al.)
**Input:** Pairwise Euclidean distance *D* in embedded 3-D space
**Output:** the pairwise MaxStep distance *D’
D’* = *D*
Let *n* be the number of data points
for *k* from 1 to *n* do
     for *i* from 1 to *n* — 1 do
          for *j* from i + 1 to n do
                *D’*(*i,j*) = min (*D’*(*i, J*) max (*D*’*i, k*),*D*’(*k, j*)))
          end
     end
end
return *D’*
~~~

~~~
**Algorithm 2:** Calculation of the Dunn index (Dunn) (adopted from Habib et al.)
**Input:** Pairwise Euclidean distance in embedded 3-D space (*D*), cluster assignment (*C*)
**Output:** Dunn index (θ)
*C_uniq* = unique(*C*)
*n* = |*C_uniq*|
Let Δ_*in*_ be an empty array of length *n*
Let Δ_*between*_ be an empty matrix of size n×n
for *i* from 1 to *n* do
            Let *ij* be the index of data whose cluster id is *C_uniq* (*i*)
            Δ_*in*_(*i*)= *max*(MaxStep(*D*(*ii*, *ii*)))
end
for *i* from 1 to *n*-1 do
            for *j* from i + 1 to *n* do
                      Let *ii* be the index of data whose cluster id is *C_uniq* (*i*) or *C_uniq* (*j*)
                      Δ_*between*_(*i, j*) = max(*MaxStep*(*D*(*ii*, *ii*)))
            end
end
*θ*= min(Δ_*between*_)/max(Δ_*in*_)
return *θ*
~~~

~~~
**Algorithm 3:** Cluster assignment
**Input:** local density (p) and local distance (5) for every cell; Pairwise Euclidean distance in embedded 3-D space (*D*).
**Output:** cluster assignment (*C*)
Let *n* be the total number of cells
Let *C*_*best*_ be an empty array of length *n*
Let *Dunn_bes_*=-1NF
for *ρ*_0_ from 0 to max (ρ) do
               for *δ*_0_ from 0 to max (δ) do
                       choose cells whose *ρ*(*i*) and *δ*(*i*) is greater than *ρ*_0_, *δ*_0_ as *Centers
                       C* = cluster_assignment(*D*, *Centers*)^*^
                       if *Dunn*(*D*, *C*) > *Dunn_best_* do
                             *Dunn_best_*=*Dunn*(*D*, *C*)
                             *C_best_* = *C*
                        end
               end
end
return *C_btst_*
*cluster assignment(*D*, *Centers*) is as described here [16]
~~~
7. **“Shuffled” cells** Due to the limited genome coverage of each single cell, cells may cluster based on their sequencing depth rather than ‘true’ co-variation^5^. To verify that our cluster results are not driven by such artefacts, we compared our results to a simulated data set. For this data set in which binary accessible sites within each cell were randomly shuffled across all accessible sites. In other words, we shuffled the data and removed the biological clustering, but maintained the distribution of sequencing depth across cells. “Shuffled” cells were uniformly distributed as a “ball” in the embedded 3-D space without clear partition of cells. However, we did observe that one of the directions becomes correlated with sequencing depth (Pearson correlation 0.55 for t-SNE3) and there is a small portion of cells that tend to form a cluster but did not pass the cut-off (*ρ*_0_,*δ*_0_) used for the P56 forebrain data set.

### Identification of cluster-specific features

We next developed a computational method which combines stability selection with LASSO^56^ to identify genomic elements (features) that potentially distinguish cells belonging to different clusters. LASSO regression enables sparse feature selections through the use of L1 penalty. However, LASSO regression often does not result in a robust set of selected features and is sensitive to data perturbation. This is especially true when features are correlated as the case here. To overcome these limitations, we adopted stable lasso to robustly identify features that distinguish every two cell clusters (Algorithm 4)^56^. Finally, we combined all identified features that distinguish different cell types.

~~~
**Algorithm 4:** Cluster specific features selection
**Input**: *X* ∈ *R*^(*n, p*)^ (*binary matrix*), *Y* ∈ {0,1}^*n*^ (*cluster label*),
*α*(*subsampling rate*), *β*(*perturbation rate*), *T*(*iteration*)
**Output**:importance score for each feature
fot *t* = 1 to *T* do
                  Randomly perturb the data:
                      Draw a subset (*X_t_, Y_t_*) of α of (X, Y)
                      Draw a vector *W*∼*U*[β,1]^p^)
                      Re-weight the features: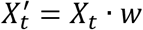
                  Compute the LASSO path of length *α*▪ *n*
                  Keep the selection matrix S_t_ ∈ ^3p, α▪n^ where ix
*S_t_*(*i, j*) = {1, *if the ith feature selected at jth step* 0, *otherwise*
end for
Compute the feature importance ix
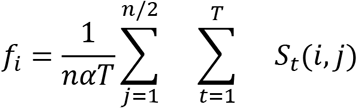
~~~

### Bulk ATAC-seq

Paired-end sequencing reads were aligned to the mm10 reference genome using Bowtie2 in paired-end mode with following parameters “bowtie2-p 5-t-X2000 ‒‒no-mixed ‒‒no-discordant^51^ and PCR duplicates were removed using SAMtools^52^. Next, mitochondrial reads were removed and the position of alignments adjusted^53^. For visualization the *bamCoverage* utility from deepTools2 was used^57^.

### Hierarchical clustering of ATAC-seq profiles in adult forebrain

DeepTools2 was used for correlation analysis and hierarchical clustering of ATAC-seq profiles from cell clusters and sorted cell-types in the adult forebrain^57^. First, we computed read coverage for each data set against the merged list of genomic elements that separate two cell clusters in the adult forebrain using the *multiBamSummary* utility. Next we used *plotCorrelation* to generate hierarchical clustering using Spearman correlation coefficient between two clusters^57^.

### Accessibility analysis and clustering of genomic elements

To cluster genomic elements based on their accessibility profile we used these promoter distal elements that were capable to distinguish two cell clusters. For each feature we extended the summits identified by MACS2^58^ in both directions by 250 bp and generated a union set of elements using *mergeBED* functionality of BEDTools v2.17.0^55^. Next, we intersected cluster specific bam files with the peak list using the *coverageBED* functionality of BEDTools v2.17.0^55^. We discarded elements that had less than five reads on average. After adding a pseudocount of one we calculated cluster-specific RPM (reads per million sequenced reads) values for each genomic element. We divided the RPM value for a given cluster by the average value of all clusters (fold over mean) and finally log2 transformed the data. The generated matrix was used for k-means clustering of the elements using Ward’s method. We performed this analysis for all adult clusters, the excitatory neuron clusters and the 12 developmental cell clusters, respectively. A list of elements for each analysis can be found in Supplementary Table 1.

### Motif enrichment analysis

To identify potential regulators of chromatin accessibility we performed motif analysis using the AME utility of the MEME suite^59^. A P-Value cut-off of < 10-5 was chosen for known motifs from the JASPAR database (JASPAR_CORE_2016_vertebrates.meme)^60^. For identification of de novo motifs HOMER tools was used with default settings^61^.

### Annotation of genomic elements

The GREAT algorithm was used to annotate distal genomic elements using following settings to define the regulatory region of a gene: Basal+extension (constitutive 1 kb upstream and 0.1 kb downstream, up to 500 kb max extension)^31^. Gene ontology categories “Molecular Function” and “Biological Processes” were used.

### Analysis of dynamic chromatin accessibility within a cell cluster

First, the ATAC-seq reads were counted in all peaks for each stage, cell type and replicate. For each cell cluster, only stages with more than 250,000 reads overlapping ATAC-seq peaks and more than 50 nuclei were used for dynamic analysis. Peaks with greater than 1 read per million reads (RPM) in at least 2 samples were kept. We used edgeR^62^ to assess the significance of difference between adjacent stages for cell clusters with at least 4 out of 7 stages passing filtering criteria. P-values were corrected using the Bonferroni method. Peaks with a Bonferroni p-value less than 0.05 were called dynamic peaks. The total number of dynamic peaks in each cell type are listed in (Extended Data Fig. 11c). For each cell type, the read counts in each peak were normalized into a unit vector (i.e values were divided by the square root of the sum of the squares of the values). K-means was used for clustering of cell clusters with more than 200 dynamic elements (K=3). Motif enrichment analysis was performed for each peak cluster using HOMER^61^.

### VISTA analysis

Genomic locations of 484 VISTA validated elements^44^ were downloaded from https://enhancer.lbl.gov using the search term “forebrain”. Genomic locations were converted from mm9 to mm10 using the *liftOver* tool (minimum rematch ratio of 0.95)^63^. 91 of these were showed specific activity in the subpallium^45^. To identify developmental clusters that are enriched for subpallial enhancers we first calculated the ratio of elements per k-means cluster overlapping with the total forebrain enhancer list and the subpallial subset separately. Finally, we calculated the relative enrichment using the ratio of subpallial over the complete forebrain regions.

### External data sets

Published ATAC-seq data of sorted excitatory neurons (GSM1541964, GSM1541965)^8^, GABAergic neurons (GSM2333635, GSM2333636)^9^, microglia (GSM2104286)^17^, neurons of the dentate gyrus (GSM2179990, GSM2179991)^27^ and distinct cortical layers (Layer2/3: GSM2333632, GSM2333633; Layer 4: GSM2333644, GSM2333645; Layer 5: GSM2333641, GSM2333642, Layer 6, GSM2333638, GSM2333639)^9^ were reprocessed. In addition, bulk ATAC-seq data for embryonic time points were analysed for comparison (https://www.encodeproject.org/search/?searchTerm=atac+forebrain, Zhao et al. manuscript in preparation)

### Data availability

Raw and processed data have been deposited to NCBI Gene Expression Omnibus with the accession number GSE1000333.

